# MYCN-induced lineage infidelity initiates SHH medulloblastoma-like tumors outside the canonical lineage of origin

**DOI:** 10.64898/2026.08.07.743303

**Authors:** Lauranne Bouteille, William Fargues, Paul de Boissier, Guillaume Girard, Mehdi Saadaoui, Cédric Maurange

## Abstract

Amplification and high expression of MYCN and MYC are recurrent features of medulloblastoma and other pediatric cancers, yet how these proto-oncogenes interact with developmental programs to initiate tumorigenesis remains unclear. Here, we combine the experimental accessibility of the avian embryo with single-cell transcriptomics to investigate how MYCN overexpression reshapes cerebellar lineage trajectories. While MYCN broadly promotes transient overproliferation, enhanced biosynthesis and delayed neural maturation, we find that these effects are insufficient to drive tumorigenesis. Instead, tumorigenic competence is restricted to a discrete population of ATOH1^+^ isthmic progenitors, which reproducibly expands into extracerebellar tumors that transcriptionally resemble SHH medulloblastoma, partially recapitulating the granule cell lineage hierarchy. By reconstructing the earliest stages of tumor initiation, we show that MYCN first expands these progenitors and then redirects them toward a hybrid PAX6⁺ granule cell progenitor-like state. Blocking PAX6 transcriptional activity abolishes tumorigenesis of the reprogrammed population, demonstrating that cooption of this lineage program is functionally required for tumor growth. Our findings reveal that MYCN-induced oncogenicity in the cerebellum proceeds through a three-step temporal sequence driven by the erosion of lineage restrictions.

## Introduction

Developmental programs generate neuronal diversity in the vertebrate brain through tightly controlled lineage trajectories that coordinate progenitor proliferation, fate specification, migration, and differentiation^1–3^. Disruption of these processes is increasingly recognized as a central mechanism underlying pediatric brain tumors, which often arise during periods of active neurogenesis and can progress rapidly ^4–6^. However, the mechanisms governing their initiation remain poorly understood. Although oncogene misexpression is a recurrent feature of pediatric cancers, how these oncogenes interact with developmental programs to drive tumor formation is unclear. In particular, it remains unclear whether they merely expand pre-existing progenitor populations and stall differentiation, or actively redirect developmental trajectories toward tumorigenic states. The relative contribution of each of these effects to tumor initiation and progression is also unknown. Defining how oncogene activation reshapes developmental trajectories is therefore essential for understanding the origins of childhood brain tumors.

Medulloblastomas are the most common malignant pediatric brain tumors ^5,7,8^. They arise in four subgroups, WNT, Sonic Hedgehog (SHH), Group 3, and Group 4, that differ in genetics, transcriptional programs, developmental features, and clinical behavior ^9–12^. Numerous lineage tracing studies and emerging developmental atlases of the mammalian cerebellum have provided high-resolution maps of cerebellar lineage trajectories^13–18^, germinal zones and progenitor states ^13,19–25^ (Figure 1A,B). Recent single-cell studies have enabled alignment of medulloblastoma cell states and trajectories with their developmental counterparts, revealing that tumor cells aberrantly redeploy fetal programs associated with distinct cerebellar lineages. This transcriptional resemblance is generally interpreted as reflecting the tumor’s cell of origin ^5,26–29^. In some cases, such predictions are supported by experimental evidence. For example, genetic manipulation of the SHH pathway in mice indicates that SHH medulloblastomas commonly arise from granule cell progenitors (GCPs) in the external granular layer (EGL) ^20,30,31^. However, the precise cells of origin of several medulloblastoma subgroups remain actively debated ^5,8^, and direct experimental evidence is largely lacking because of the difficulty of manipulating and tracking cerebellar progenitors with precise spatial and temporal control.

**Figure 1:**
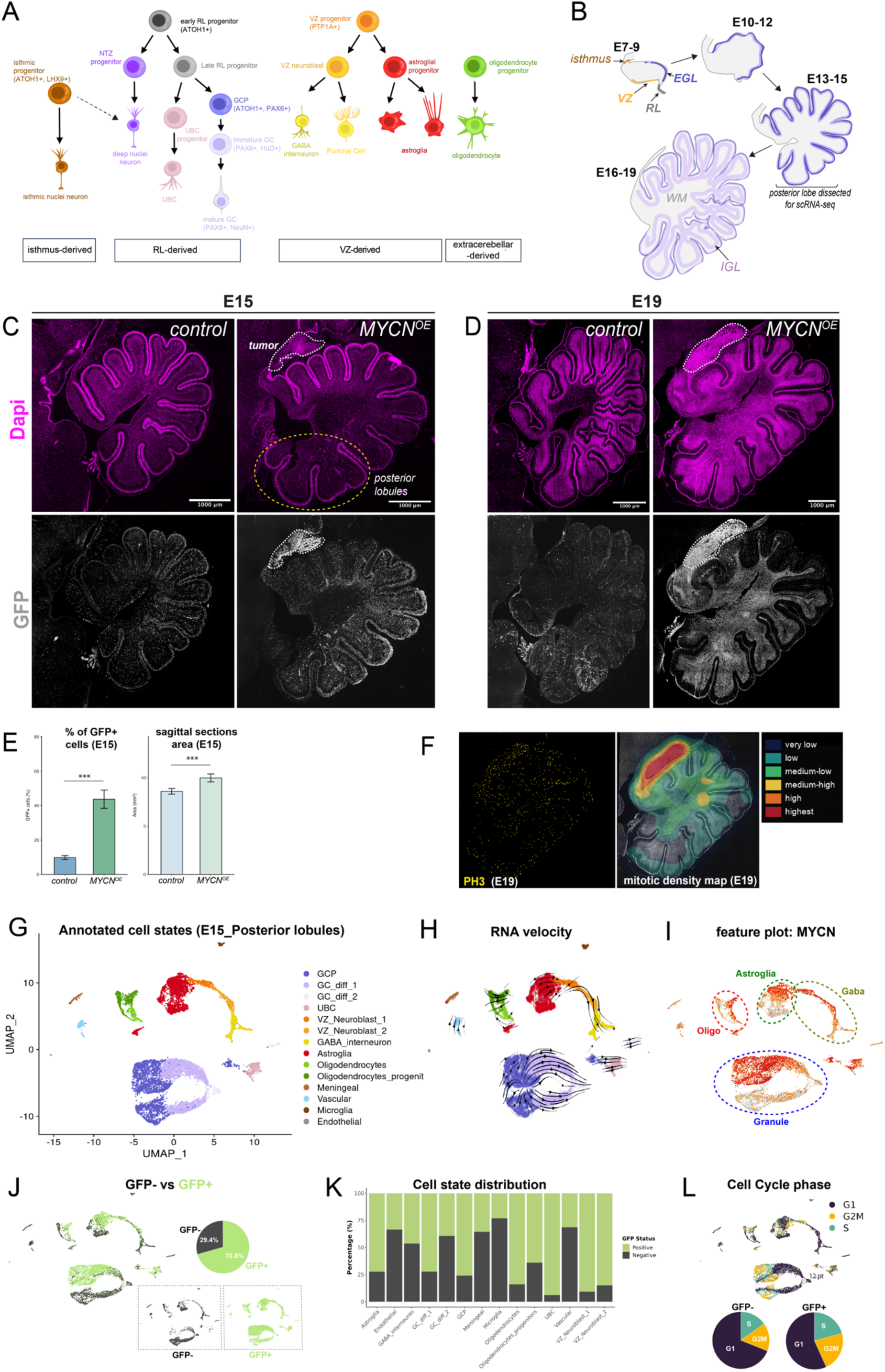
Most cerebellar lineages are resilient to the oncogenic effects on MYCN. (A) Schematics of cerebellar lineages and (B) Schematic sagittal sections of the developing chick cerebellum at successive embryonic stages, highlighting the main proliferative centers: the rhombic lip (RL), ventricular zone (VZ), external granular layer (EGL), and isthmus. Cerebellar neurons arise from these four spatially distinct progenitor pools in distinct temporal waves ^13,46^. In particular, glutamatergic neurons derive from ATOH1⁺ progenitors of the rhombic lip (RL), which first produce progenitors of deep cerebellar nuclei neurons, and then give rise to unipolar brush cell (UBC) progenitors and granule cell progenitors (GCPs) ^22,23,47^. The latter migrate out of the rhombic lip and expand extensively to form the EGL before differentiating into granule neurons (also known as granule cells (GCs) ^20,25,30^. Mature GCs then migrate to the internal granule layer (IGL), composing the largest neuronal population of the brain. In parallel, Purkinje cells and then GABAergic neurons arise from the PTF1A⁺ ventricular zone (VZ), together with astroglial cell types ^19,48^. In addition, a third, less well-described, progenitor zone localizes in the isthmic region, at the border between the cerebellum and the midbrain, and produces a distinct set of isthmic nuclei neurons, possibly also contributing to deep cerebellar nuclei ^16,18,21,49^. The outer EGL, containing granule cell progenitors (GCPs), is shown in dark purple, whereas granule cells (GCs) migrating toward the internal granular layer (IGL) are shown in light purple. The white matter (WM) is depicted in light gray. (C and D) Immunofluorescence images of sagittal cerebellar sections from control and MYCN-overexpressing (MYCN-OE) embryos at 13 days post electroporation (13 DPE) (C) and 17 DPE (D) corresponding to E15 and E19 respectively. Sections are stained for GFP (green) to label cells deriving from electroporated progenitors and DAPI (blue) to visualize nuclei. White dotted outlines indicate an extracerebellar hyperplasia located at the isthmic region in MYCN-OE cerebella. Yellow dashed outline indicates the posterior lobules dissected for single-cell RNA-seq (See Figure 2). (E) Quantification of the percentage of GFP⁺ cells (left) and of cerebellar section area (right) in control and MYCN-OE embryos at 13 DPE. Data are represented as mean ± SEM. Results are representative of at least 3 sections per embryo and a minimum of 3 embryos per condition. Statistical comparisons were performed using the Wilcoxon rank-sum test. *p < 0.05, **p < 0.01, ***p < 0.001, ns = not significant. (F) Mitotic density map of a sagittal cerebellar section at E19 generated from an immunostaining image labeling phospho-Histone 3 (PH3) and overlaid onto the corresponding DAPI-stained cerebellar section images. (G) UMAP projection of single-cell RNA sequencing data from the MYCN-OE posterior cerebellar lobe showing annotated cell states, including granule cell progenitors (GCP), differentiating granule cells (GC_diff_1 and GC_diff_2), unipolar brush cells (UBC), ventricular zone neuroblasts (VZ_Neuroblast_1 and VZ_Neuroblast_2), GABAergic interneurons, astroglia, oligodendrocytes and oligodendrocyte progenitors, vascular cells, meningeal cells, microglia, and endothelial cells. (H) RNA velocity analysis projected onto the UMAP embedding showing inferred developmental trajectories across cerebellar cell populations. Streamlines indicate predicted transcriptional dynamics and lineage progression. (I) Feature plot depicting MYCN expression after electroporation in the various lineages. (J) UMAP projection highlighting GFP− and GFP+ cells with accompanying pie charts showing the relative proportion of GFP− and GFP+ cells in the dataset. (K) Proportions of GFP^+^ and GFP^-^ cells obtained from the single-cell RNA-seq of posterior lobules, according to cell types. Note that GFP+ cells are predominant in progenitors such as GCPs and VZ-neuroblasts, but minor in more differentiated cell states like GC_diff_2 and GABA_interneurons. (L) Cell cycle phase distribution across the UMAP embedding and comparison of cell cycle states between GFP− and GFP+ populations. Pie charts indicate the proportion of cells in G1, S, and G2/M phases for each population.

MYC-family transcription factors (MYC, MYCN and MYCL) are central regulators of proliferation, stem cell maintenance and metabolism, with MYCN more specifically expressed in the nervous system during early development ^32–34^. In the developing cerebellum, MYCN is a target of SHH in GCPs in the EGL and is required for GC amplification during development ^32,33,35,36^. MYC and MYCN have a well-known oncogenic activity, and their amplification or overexpression is associated with a broad range of cancers and poor prognosis. In medulloblastoma, MYC is particularly associated with aggressive Group 3 tumors, whereas MYCN is commonly linked to the SHH subgroup; Group 4 tumors may overexpress either gene ^11,37–39^. Elevated levels of MYC or MYCN expression is associated with poor clinical outcome. Studies investigating the oncogenic effects of MYCN overexpression in cerebellar progenitors have reported divergent outcomes depending on the experimental context. In both genetically engineered mouse models (GEMMs) and graft-based models using in vitro–differentiated human cerebellar progenitors, MYCN overexpression with or without p53 inactivation gave rise to tumors resembling either Group 3 or SHH medulloblastoma ^34,40–45^. Moreover, these approaches did not conclusively identify the cell of origin or the mechanisms driving transformation. Addressing this question requires experimental systems that permit oncogene activation during early neural development while preserving the spatial and temporal context needed to follow lineage behavior *in vivo*.

Here, we exploit the accessibility of the early chick embryo to mosaically overexpress MYCN in the various neuroepithelial progenitors of the midbrain-hindbrain region and examine how oncogenic activation reshapes cerebellar developmental trajectories. Combining single-cell transcriptomics, lineage analysis, and developmental imaging, we find that MYCN overexpression activates a shared biosynthetic program that promotes proliferation and impairs differentiation across cerebellar lineages, but is not sufficient to induce tumorigenesis. Surprisingly, we consistently observed at the isthmus the formation of an extracerebellar tumor resembling SHH medulloblastoma. Unexpectedly, tumor progenitors displayed a hybrid transcriptional identity combining GCP and isthmic progenitor markers. Developmental analyses reveal that MYCN reprograms isthmic progenitors toward a GCP-like state several days after the onset of overexpression, and that acquisition of this hybrid identity is required for transition to tumorigenesis, likely through the co-option of PAX6-dependent transcriptional programs. Together, our findings demonstrate that oncogene-driven lineage infidelity is a key driver of tumor initiation in the developing cerebellum, challenging the notion that medulloblastomas mirror their lineage of origin.

## Results

### The developing cerebellum is globally resilient to MYCN except at the isthmus

To investigate how MYCN misexpression affects cerebellar development (Figures 1A and 1B), we took advantage of the accessibility of the chick embryo at embryonic day 1.5 (E1.5) to electroporate neuroepithelial cells in the midbrain–hindbrain region with a multicistronic plasmid driving co-expression of GFP and chick MYCN. Genomic integration enabled sustained transgene expression in electroporated cells and their progeny. Notably, cerebellar development is accelerated in avian species relative to mammals, enabling lineage-specific differentiation defects to be assessed before birth ^25^. At E15, corresponding to 13 days post-electroporation (13 DPE), embryos electroporated with a control GFP-only plasmid displayed a sparse distribution of GFP^+^ cells throughout the cerebellum, including the EGL, IGL and white matter (WM) (Figure 1C). This pattern indicates that multiple cerebellar progenitor populations were targeted and that their progeny contributed broadly to cerebellar structures. GFP^+^ cells were also detected outside the cerebellum (Figure 1C), consistent with the fact that electroporation was not restricted to cerebellar progenitors.

In contrast, co-electroporation with MYCN consistently resulted in a marked increase in the number of GFP^+^ cells throughout the cerebellum, suggestive of enhanced proliferation (Figures 1C-1E). Consistently, cerebella overexpressing MYCN exhibited an increased volume (Figures 1E). In some anterior lobules, the EGL contained comparatively fewer GFP^+^ cells, whereas highly electroporated lobules appeared mildly distorted, likely reflecting excessive proliferation in the EGL and/or WM. The choroid plexus was frequently strongly electroporated, yet never displayed overt overgrowth, indicating that not all cell types or tissues respond to MYCN with hyperproliferation. At E19 (17 DPE), the overall laminar architecture of the cerebellum remained largely preserved, arguing against tumor-like overgrowth within the cerebellum itself (Figure 1D).

By contrast, at both E15 and E19, a prominent hyperplastic mass was observed anterior to the cerebellum in more than 70% of electroporated cerebella (n>20), apparently emerging from a region adjacent to the isthmus, at the cerebellum–midbrain boundary (Figures 1C and 1D). This hyperplasia showed rapid growth and an increased density of mitotic cells at late embryonic states (E19) compared to other regions of the cerebellum, indicative of sustained proliferation (Figure 1F). The near- systematic appearance of this tumor-like hyperplasia argues against additional stochastic mutations as the underlying cause. Instead, these findings indicate that the response to MYCN overexpression is heterogeneous and likely lineage-dependent, with few progenitors immediately sensitive to its oncogenic activity.

To better investigate the impact of MYCN overexpression (MYCN^OE^) across cerebellar lineages, we performed single-cell RNA sequencing at E15, 13 days after electroporation of the MYCN::GFP plasmid (13DPE). We first dissected the posterior cerebellum, including the three most posterior lobules (Figure 1C). This preparation was expected to contain both electroporated and non-electroporated cells, the latter serving as an internal wild-type reference population.

Our dataset provides to our knowledge the first single-set transcriptomic atlas of a developing chick cerebellum. We used transcriptomic signatures from human cerebellar cell types for annotation ^17^, and applied RNA velocity analysis to infer lineage relationships. This approach resolved multiple clusters corresponding to canonical cerebellar developmental trajectories of this region (Figures 1G and 1H, and Table S1). The UMAP representation depicted astroglial (SOX9^+^) and GABAergic (PAX2^+^) lineages arising from a common pool of ventricular zone neuroblasts (VZ_NBs) (PTF1A^+^) (Figures 1G, 1H, S1A and S1B). The GC lineage was identified by high levels of *PAX6* and *BARHL1* expression, with GCPs coexpressing cell cycle genes (*CDC20, CDK1, TOP2A*) and *ATOH1*, while differentiating GCs expressed *RELN, EOMES*, and *NEUROD1* (Figure S1A and S1C). Other clusters included oligodendrocytes (SOX8^+^ and OLIG2^+^), possible UBCs, (LMX1B^+^, LINGO2^+^, OTX2^+^, EOMES^+^) although this population remains to be characterized in chicken ^13^ (Figures 1G, S1A, S1D and S1E). Smaller clusters were identified as endothelial, meningeal, microglial, and vascular cells (Figures 1G and S1A). Notably, Purkinje cells were not clearly identified in the single-cell dataset, despite being readily detected by immunostaining at this stage (Figures 1G and S1F), likely owing to their loss during dissociation because of their large size. Glutamatergic nuclei neurons are also absent as they are located more anteriorly than the posterior region used for single-cell RNA-seq.

Expectedly, MYCN expression strongly overlapped with GFP expression (Figures 1I and 1J). Importantly, all lineages contained both GFP^+^ and GFP^−^ cells, indicating that our electroporation strategy targeted multiple progenitor populations in a mosaic manner (Figures 1J and 1K), allowing us to study global and specific consequences of MYCN overexpression across lineages.

Single-cell RNA-seq confirmed the overrepresentation of MYCN^+^ GFP^+^ cells (71% versus 29%; Figure 1J). These MYCN^+^ GFP^+^ population also depicted an increased proportion of S-phase and G2/M-phase cells (Figure 1L). The fraction of progenitor populations, including GCPs and VZ neuroblasts, was similarly increased in the GFP^+^ compartment (Figure 1K). In contrast, the proportion of more differentiated cells such as GABAergic interneurons or differentiated GC (GC_diff_2) was decreased (Figure 1K). Together, these results support the conclusion that MYCN overexpression globally promotes enhanced proliferation without inducing tumor growth in the context of the embryonic cerebellum. In contrast, a region at the isthmus appears particularly sensitive to the oncogenic activity of MYCN.

### MYCN triggers a pan-lineage biosynthetic and maturation-delay program that is insufficient for tumor formation

To determine how MYCN overexpression affects transcriptional programs during cerebellar development, we selected the top200 upregulated and top200 downregulated genes between GFP^+^ and GFP^-^ cells for each lineage separately (astroglial, GABAergic, granule, and oligodendrocyte) and then analyzed their overlap (Figures 2A and 2B and Tables S2 and S3). This analysis revealed a two- armed transcriptional logic operating at two levels of specificity.

**Figure 2.**
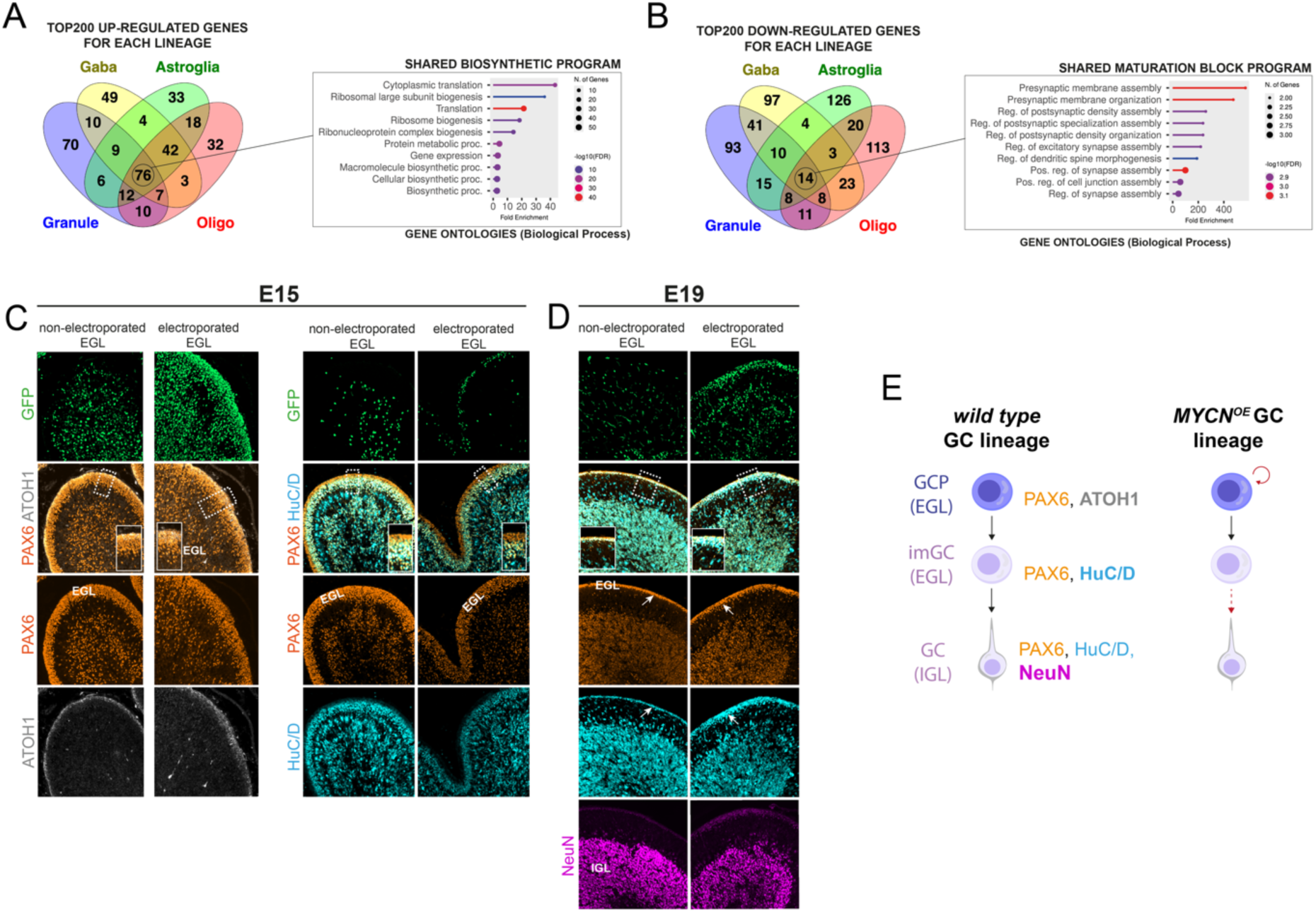
: MYCN^OE^ triggers shared and lineage-specific transcriptional changes. (A) Venn diagram depicting overlaps between the top 200 upregulated genes in the various lineages after MYCN^OE^. Gene ontology was performed for biological processes for genes upregulated in all lineages. (B) Venn diagram depicting overlaps between the top 200 downregulated genes in the various lineages after MYCN^OE^. Gene ontology was performed for biological processes for genes downregulated in all lineages. (C) Sagittal section of lobules of an electroporated cerebellum at E15. Poorly and highly electroporated lobules are shown for comparison. Lobules are labelled for GFP, PAX6 and HuC/D by immunostaining, and ATOH1 by RNA FISH. GFP^+^ cells co-express MYCN. Dashed boxes indicate part of the EGL enlarged in the lower right corner. (D) Sagittal section of lobules of an electroporated cerebellum at E19. Poorly and highly electroporated lobules are shown for comparison. Lobules are labelled for GFP, PAX6, HuC/D and NeuN by immunostaining. GFP^+^ cells co-express MYCN. Dashed boxes indicate part of the EGL enlarged in the lower right corner. Arrow indicates the reduced EGL at E19 with accumulation of HuC/D+ immature GCs in the electroporated lobule. (E) Schematic summary of granule lineage progression in control cerebellar lobules and MYCN-electroporated lobules. Red arrows show perturbations associated with MYCN overexpression (increased progenitor self-renewal, delayed differentiation. imGC: immature GC.

Among the top200 upregulated genes, a remarkably large number (76) was common to all lineages (Figure 2A). Gene Ontology analysis revealed a strong biosynthetic program dominated by genes involved in ribosome biogenesis (over 35 *RPL/RPS* genes), translation (*YBX1, EEF1B2, EEF2, EIF5A2*), nucleolar function (*NPM1, NPM3, NCL, NOLC1, NOP56*), proteostasis (*HSP90AB1, HSPD1, HSPE1*), and mitochondrial support (*TOMM20, ENO1* and *PKLR*) (Figures 2A, S2A and S2B, and Table S3). In addition, increased RNA counts in GFP^+^ cells indicated a global enhancement of transcriptional activity (Figure S2C). This shared response to MYCN overexpression is consistent with activation of a strong anabolic state across all lineages.

Conversely, MYCN broadly repressed genes associated with neuronal differentiation, adhesion, and synaptic function. Fourteen genes were consistently downregulated in all four lineages including *ADGRL3, NELL1, PTPRD, CHL1, NLGN1, LSAMP, NKAIN2* and *PCDH9*, indicating a general suppression of neuronal maturation programs (Figures 2B and S2A, and Table S3).

Superimposed on this shared program, MYCN elicited lineage-specific responses. The granule lineage exhibited the largest number of uniquely upregulated genes (70/200) (Figure 2A). Of those, *TENM4* encoding *Teneurin-4*, a large transmembrane protein involved in neuronal connectivity and synaptic organization, was not detected in the non-electroporated granule cells (Figure S2A and S2D), suggesting that MYCN, beyond boosting biological processes associated with anabolic programs, can induce ectopic gene expression *de novo*, possibly affecting cell identities. Suppression of neuronal maturation was particularly pronounced in neuronal lineages. Indeed, in granule and GABAergic cells, MYCN repressed multiple genes involved in synaptic organization and excitability, including *DLG2, HCN1, CNTNAP2, KCNQ3, GRM5, NBEA*, and *CADM2* (Figure S2A and Table S3). In addition, GABAergic cells tend to maintain an immature VZ-like neurogenic program (*NOTCH1, ASCL1, PTF1A, PAX7, SOX11, RORB, POU3F3*) (Figure S2A and Table S3) consistent with the increased fraction of VZ progenitors among GFP^+^ cells (Figure 1K). Interestingly, MYC was strongly silenced by MYCN in the granule lineage, suggesting that MYC and MYCN expression may be mutually exclusive in this lineage (Figures S2A and S2D).

The single-cell analysis indicates that, at E15 after MYCN overexpression, the granule lineage, the presumptive lineage of origin of SHH medulloblastomas, is enriched in GCPs and exhibit impaired differentiation. To validate this effect *in situ* and investigate the impact at later stages, we studied the expression of key markers of granule lineage progression at E15 and E19 with or without MYCN overexpression. PAX6 was used to label the granule lineage, ATOH1 for GCPs, HuC/D for post-mitotic GCs, and NeuN for mature GCs that had migrated into the internal granule layer (IGL). Electroporated lobules were compared with non-electroporated ones of the same cerebellar slice for internal control. At E15, the EGL of non-electroporated lobules displayed the expected two-layered organization: a thin outer layer of PAX6^+^ ATOH1^+^ GCPs and a broader inner layer of PAX6^+^ ATOH1^−^ HuC/D^+^ immature GCs (Figures 2C). In electroporated lobules, this organization was largely preserved, with distinct ATOH1^+^ and HuC/D^+^ layers still evident. NeuN staining remained largely absent from the IGL in both conditions at this stage, indicating that most GCs had not yet initiated migration (Figure S3). Thus, at E15, MYCN- OE does not appear to strongly impair the transition from GCPs to post-mitotic immature GCs within the EGL, although the ATOH1^+^ outer layer may be slightly thicker (not quantified).

By E19, in non-electroporated lobules, the immature HuC/D^+^ GC pool in the EGL was markedly reduced compared to E15, while in parallel, a prominent NeuN^+^ internal granule layer (IGL) was detected indicating that mature GCs had migrated from the EGL (Figure 2D). In the EGL, all cells were HuC/D^+^ indicating that the HuC/D^-^ ATOH1^+^ outer layer seen at E15 had disappeared (Figure 2D vs 2C). Thus, at this stage most if not all ATOH1^+^ progenitors have progressed to the immature GC state. In electroporated lobules, all PAX6^+^ cells of the EGL were likewise HuC/D^+^ indicating that progression towards the immature GC state was not blocked by MYCN^OE^ (Figure 2D). However, GCs tended to accumulate in the EGL as immature HuC/D^+^ NeuN^−^ cells suggesting delayed migration in the IGL and impaired differentiation (Figure 2D, arrow). Thus, GCPs overexpressing MYCN appear to undergo transient overproliferation, but retain the ability to exit the cell cycle and progress to a more differentiated, albeit still immature GC state, without forming tumors (Figure 2E). Given the high expression of MYCN in many SHH medulloblastomas and the proposed role of GCPs as a cell of origin, these experiments reveal an unexpected resilience of the granule cell lineage to MYCN-driven oncogenicity.

Together, these findings indicate that MYCN promotes a shared anabolic program, but also broadly restrains lineage maturation, thereby reshaping developmental progression in a lineage-dependent manner. However, although these perturbations may prime for transformation, they do not appear sufficient to drive oncogenic growth within the embryonic cerebellum.

### The MYCN-induced isthmic tumor exhibits an SHH medulloblastoma-like granule lineage hierarchy

To better understand what drives tumorigenesis from the isthmus in particular, we performed single-cell RNA-seq of the extracerebellar tumor at E15 (13DPE). The transcriptomic analysis showed at this stage that ∼90% of cells within the tumor expressed GFP, indicating that the hyperplasia is largely composed of electroporated cells overexpressing MYCN (Figure S4).

To place these cells in the context of normal cerebellar lineages, we compared the tumor dataset with single-cell profiles from the posterior cerebellum of the same embryo electroporated with MYCN, as well as a control posterior cerebellum electroporated with GFP alone. When the three datasets were merged without integration, most cells largely intermingled, except for a prominent cluster of tumor cells that remained separated (Figure 3A). This cluster represented 58% of the cells recovered from the extracerebellar tumor. Because posterior cerebellar cells from the control and MYCN^OE^ conditions mixed extensively, and because a subset of tumor cells clustered well with the GABAergic, astroglial and oligodendrocyte lineages of the other conditions, the presence of a large single segregating tumor cluster is unlikely to reflect batch effects, but rather suggests a transcriptionally distinct population absent from posterior cerebellar lobules.

**Figure 3.**
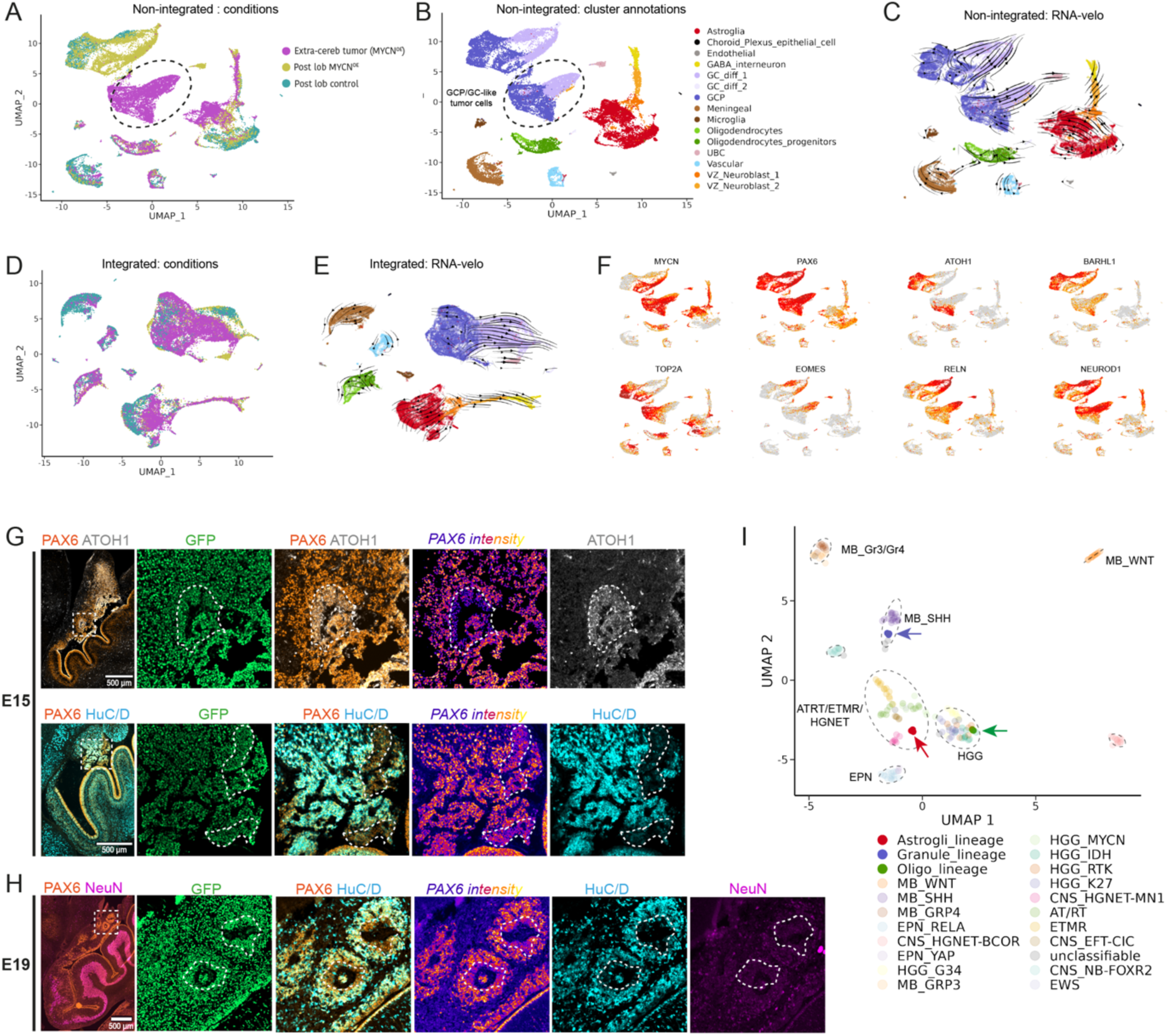
: MYCN-induced extra-cerebellar tumor cells display granule lineage identity and resemble SHH medulloblastoma. (A) UMAP of merged single-cell RNA-seq datasets from E15 samples without batch integration. Cells derive from posterior cerebellar lobules electroporated with GFP only (Post lob control), posterior lobules electroporated with GFP and MYCN (Post lob MYCN^OE^), and the extracerebellar tumor electroporated with GFP and MYCN (Extra-cereb tumor). Tumor cells form a large cluster that remains distinct from posterior cerebellar populations (dotted circle). (B) UMAP colored by signature-based cell-type annotations. The large tumor cluster is assigned to the granule cell lineage, including granule cell progenitor (GCP) and differentiating granule cell (GC) states. (C) RNA velocity projected onto the non-integrated UMAP reveals a trajectory consistent with GCP-to-GC differentiation within the tumor cluster. (D) UMAP after anchor-based integration of the three datasets, colored by condition. Granule lineage-like tumor cells merge with granule lineage populations from the posterior cerebellum. (E) RNA velocity projected onto the integrated UMAP confirms preservation of developmental trajectories across granule lineage populations. (F) Feature plots showing expression of representative granule lineage markers. Tumor cells express *MYCN*, the granule lineage markers *PAX6* and *BARHL1*, GCP markers (*ATOH1, TOP2A*), and markers of differentiating granule cells (*EOMES, RELN, NEUROD1*). (G-H) The left panels show images at the cerebellar/midbrain boundary. The dashed box represents the region of the extracerebellar tumor that is enlarged on the right panels. Most cells of the tumor are GFP^+^. (G) At E15, the tumor contains a disorganized PAX6^+^ cell population with a subset of *ATOH1^+^* cells expressing lower levels of PAX6 (dashed outline). PAX6 immunostaining is color-coded as a function of its intensity. More differentiated HuC/D^+^ GC-like cells express higher levels of PAX6. ATOH1 expression is assessed by RNA-FISH. (H) At E19, the PAX6low HuC/D-GCP-like population persist. NeuN is absent in the tumor indicating an absence of fully differentiated GC-like cells. (I) UMAP of pseudo-bulk transcriptomic profiles comparing extracerebellar tumor-derived cell populations with reference datasets from human pediatric brain tumors ^50^. The dominant tumor GC-like population (blue arrow) clusters with SHH medulloblastoma (MB_SHH), whereas astroglial-like (red arrow) and oligodendrocyte-like (green arrow) tumor populations group with neuroepithelial tumors and high-grade gliomas, respectively.

Despite this segregation, annotations based on cerebellar lineage signatures assigned the large tumor cluster to the granule cell lineage (Figure 3B). RNA velocity analysis revealed a coherent differentiation trajectory from annotated GCPs to GCs (Figure 3C). In agreement with this granular-like identity, anchor-based integration across the three datasets resulted in the merging of tumor granule-like clusters with granule cell lineage clusters from the posterior cerebellum (Figures 3D and 3E). Consistently, tumor cells of this large cluster expressed canonical granule lineage markers, including *PAX6* and *BARHL1*, with *ATOH1* and cell-cycle genes such as *TOP2A* enriched in GCP-like subclusters, and *EOMES, RELN, NEUROD1*, and *NEUROD2* marking differentiating GC-like cells (Figure 3F).

We confirmed by immunostaining and RNA-FISH the existence of a PAX6^+^ ATOH1^+^ GCP-like subpopulation at E15 in the tumor, intermingled with more differentiated HuC/D^+^ cells expressing higher levels of PAX6 (Figure 3G). The PAX6^low^ HuC/D^-^ GCP-like population was maintained in the tumor at E19 showing sustained self-renewal and amplification properties beyond normal GCPs in the EGL (Figure 3H). NeuN was absent from the tumor indicating that mature GCs failed to be produced (Figure 3H). Thus, MYCN overexpression can generate an ectopic tumor composed largely of granule lineage- like cells in a region adjacent to the isthmus.

To further assess the relationship between this tumor and human pediatric brain tumors, we generated pseudo-bulk transcriptomes from the three major cell types present in the extracerebellar tumor and compared them with reference transcriptomic datasets from human brain tumor cohorts^50^. Principal component and clustering analyses showed that the dominant granule-like tumor population clustered with Sonic Hedgehog (SHH) medulloblastoma, whereas the smaller astroglial-like and oligodendrocyte-like populations grouped with neuroepithelial tumors and high-grade gliomas respectively (Figure 3I). These results indicate that the MYCN-induced extracerebellar tumor is predominantly composed of cells from a granule-like lineage that shares transcriptional and hierarchical features with human SHH medulloblastoma.

### The MYCN-induced extracerebellar tumor contains progenitors with a mixed granule/isthmic lineage identity

Although co-expression of PAX6 and ATOH1 is a defining feature of GCPs, we were intrigued by the observation that tumor GCP-like cells did not merge with posterior lobe GCPs in the non-integrated single-cell analysis (Figures 3A and 3D). This suggested that, despite their apparent granular identity, GCP-like cells in the tumor differ substantially from GCPs located in the EGL.

To define these differences and understand how GCP-like progenitors become tumorigenic, we used our single-cell RNA-seq data to identify the top 100 differentially-expressed genes defining the two cell types (Table S4). Consistent with our observation that differentiation is stalled in the hyperplasia, genes associated with neuronal maturation and circuit formation were downregulated in tumorigenic progenitors (Figure 4A). In addition, this was accompanied by a marked difference in metabolic gene expression between the two cell types. Tumor progenitors downregulated genes involved in serine biosynthesis and amino acid metabolism whereas genes associated with glycolysis (*LDHA*), the pentose phosphate pathway (*RPIA*), and biosynthetic growth (*RPL22L1*) where highly up-regulated (Figure 4A). Thus GCP-like tumor progenitors exhibit a distinct metabolic profile than MYCN^+^ GCPs in the EGL.

**Figure 4.**
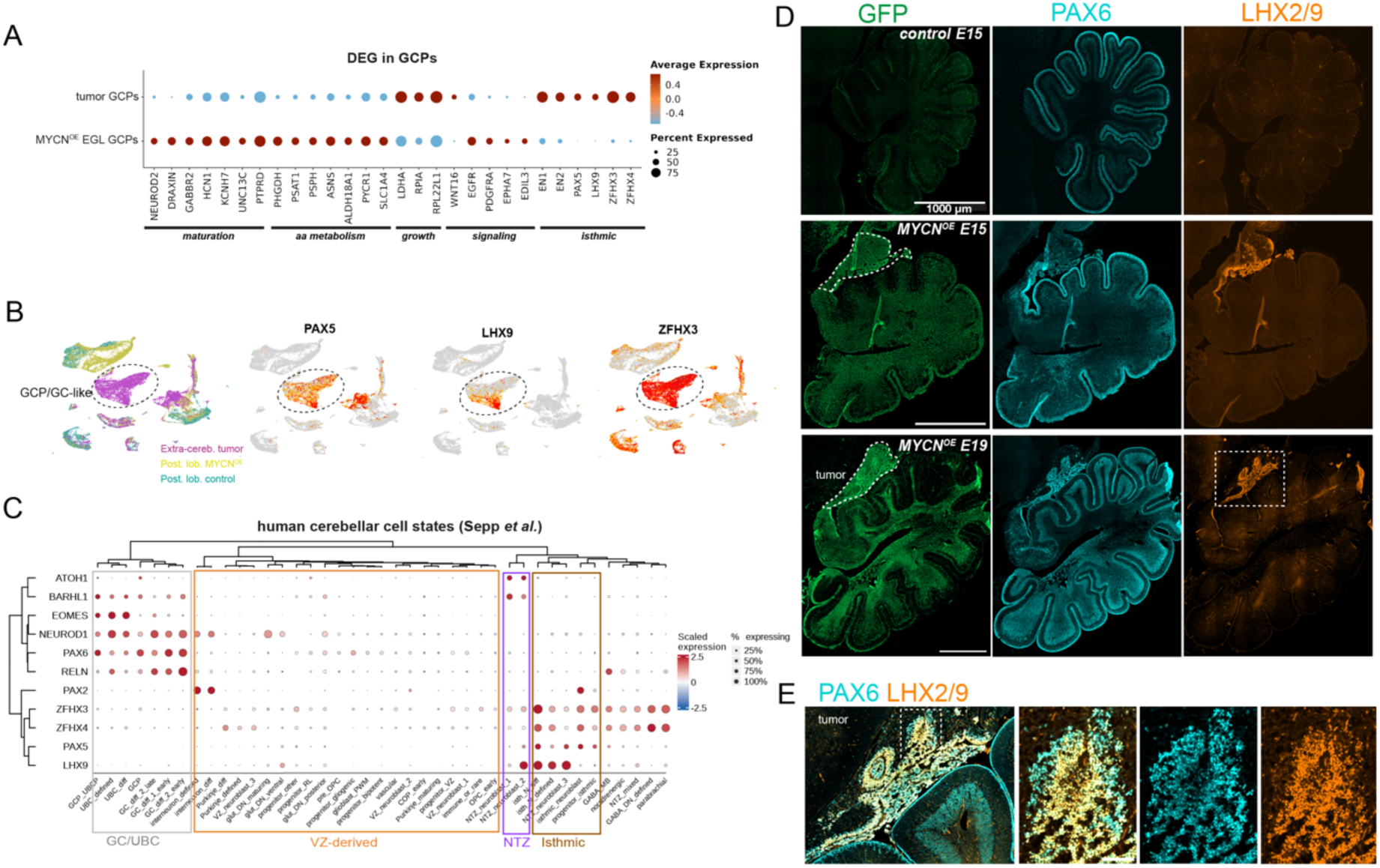
MYCN-induced tumor cells display a hybrid granule/isthmic lineage identity. (A) Dot plot showing a set of top differentially expressed genes in GCPs overexpressing MYCN from the EGL or tumor. (B) UMAP representation of merged single-cell RNA-seq datasets from control posterior cerebellum, MYCN-electroporated posterior cerebellum, and the extracerebellar tumor. Tumor GC-like lineage is highlighted by the dotted line. Feature plots show expression of *PAX5*, *LHX9* and *ZFHX3* highlighting their enrichment in tumor cells. (C) Dot plot showing expression of representative transcription factors across cerebellar and brainstem lineages in human. *LHX9, ZFHX3, ZFHX4* and *PAX5*, characteristic of isthmic nuclei neurons, are never co-expressed with granule lineage markers (*PAX6, BARHL1, ATOH1*) during normal cerebellar development. (D) Sagittal sections showing GFP, PAX6, and LHX2/9 immunostaining at E15 and E19. LHX2/9 expression is absent from control cerebellar tissue but strongly expressed in the extracerebellar tumor following MYCN overexpression. (E) Higher-magnification images of the tumor at E19 showing that LHX2/9 is expressed in a subset of PAX6^+^ tumor cells.

In parallel, tumor progenitors showed differential expression of various genes involved in developmental signaling pathways. For example, *WNT16* was strongly up-regulated while *EGFR, PDGFRA, EPHA7*, as well as the integrin ligand *EDIL3* were downregulated (Figure 4A). This suggests that normal GCPs and tumorigenic GCP-like cells evolve in distinct niches, responding to distinct microenvironmental signals.

Strikingly, in addition to these differences, tumor progenitors displayed high expression levels of *EN1, EN2, PAX5, LHX9, ZFHX3* and *ZFHX4* (Figures 4A), which are characteristic of isthmic nuclei neuronal signatures in mammals ^16–18,21,51^. *PAX5, ZFHX3* and *ZFHX4* were broadly expressed throughout the tumor population, whereas *LHX9* expression was restricted to the *ATOH1^+^* GCP-like compartment (Figures 4B and 3F). To investigate whether this mixed signature displaying granule and isthmic features reflects an existing cell population during development, we turned to recently generated single-cell data produced from developing human cerebellum ^17^. Expectedly, typical markers of granule and isthmic lineages, such as those present in the extracerebellar tumor, clustered separately according to their expression pattern in the various cell types. Importantly, no single cell type showed co-expression of all these markers, possibly implying that this phenomenon could be a tumor-specific feature reflecting an abnormal hybrid developmental identity adopted by MYCN-overexpressing tumor progenitors (Figure 4C). Alternatively, this could reflect the amplification of an avian specific progenitor population.

We then turned to cerebellar slices to validate these data in situ. We used an antibody recognizing both LHX9 and LHX2. Immunostaining did not detect LHX2/9^+^ cells in the avian cerebellum or isthmic region at E15 under control conditions, whereas strong LHX2/9 expression was observed in the tumor at both E15 and E19 following MYCN overexpression (Figure 4D). Higher-magnification analysis showed that LHX2/9 expression was confined to the PAX6^low^ population, consistent with its association with the ATOH1^+^ GCP-like compartment (Figure 4E; see also Figure 3G). Notably, this LHX2/9^+^ GCP-like population (hereafter referred to as LHX9^+^) remained proliferative at E19, suggesting that it contributes to sustained tumor growth (Figure S5A).

In addition, we detected groups of cells co-expressing GFP, PAX6 and LHX9 within the EGL of the most anterior lobules, adjacent to the tumor, indicating that tumor progenitors may potentially migrate and preferentially invade the endogenous GCP niche, consistent with a tropism conferred by their GCP-like identity (Figure S5B).

Together, these findings indicate that the MYCN-induced extracerebellar tumor is sustained by a GCP-like progenitor population endowed with a peculiar metabolism and signaling niche, possibly invasive properties, and a hybrid granule/isthmic identity.

### MYCN progressively reprograms an early LHX9^+^ isthmic progenitor population towards a mixed PAX6^+^ GCP-like fate

The expression of transcription factors characteristic of isthmic nuclei neurons in tumors suggested that the extracerebellar tumor may originate from an early progenitor population located near the cerebellar isthmus. We therefore sought to trace tumor initiation back to a potential LHX9^+^ PAX6^+^ ATOH1^+^ cell-of-origin during earlier stages of cerebellar development.

In mammals, most LHX9^+^ neurons are generated in the developing cerebellum before GCPs to form the lateral cerebellar nucleus (also known as dentate nucleus). They mostly arise from ATOH1^+^ early rhombic lip progenitors although a subset of them may originate from an ATOH1^+^ progenitor pool located at the isthmus largely responsible for generating isthmic nuclei. The lateral cerebellar nucleus is absent in birds but the isthmic ATOH1^+^ LHX9^+^ progenitor pool is known to be more prominent, leading to the formation of an additional bird-specific isthmic nucleus ^16,21,49,52^. Consistent with this model, we detected a small population of ATOH1^+^ LHX9^+^ cells in the isthmus at E9 in control embryos, while ATOH1^+^ GCPs distributed along the EGL lacked LHX9 expression (Figure 5A). Notably, PAX6 expression was restricted to GCPs within the EGL and was absent from the ATOH1^+^ LHX9^+^ isthmic domain at this early stage (Figure 5A). Both progenitor populations were efficiently targeted by electroporation, as indicated by dispersed GFP^+^ cells at E9 (7DPE) (Figure 5A). In the isthmic region, about 20% of LHX9^+^ domain was GFP^+^ (Figure 5B). Upon MYCN overexpression, clusters of GFP^+^ cells represented more than 50% of the total LHX9^+^ isthmic domain at E9, consistent with the ability of MYCN to boost progenitor proliferation through activation of a biosynthetic program (Figure 5B and 5C). However, electroporated isthmic progenitors did not express PAX6 at this stage like their wild type counterparts (Figure 5C).

**Figure 5.**
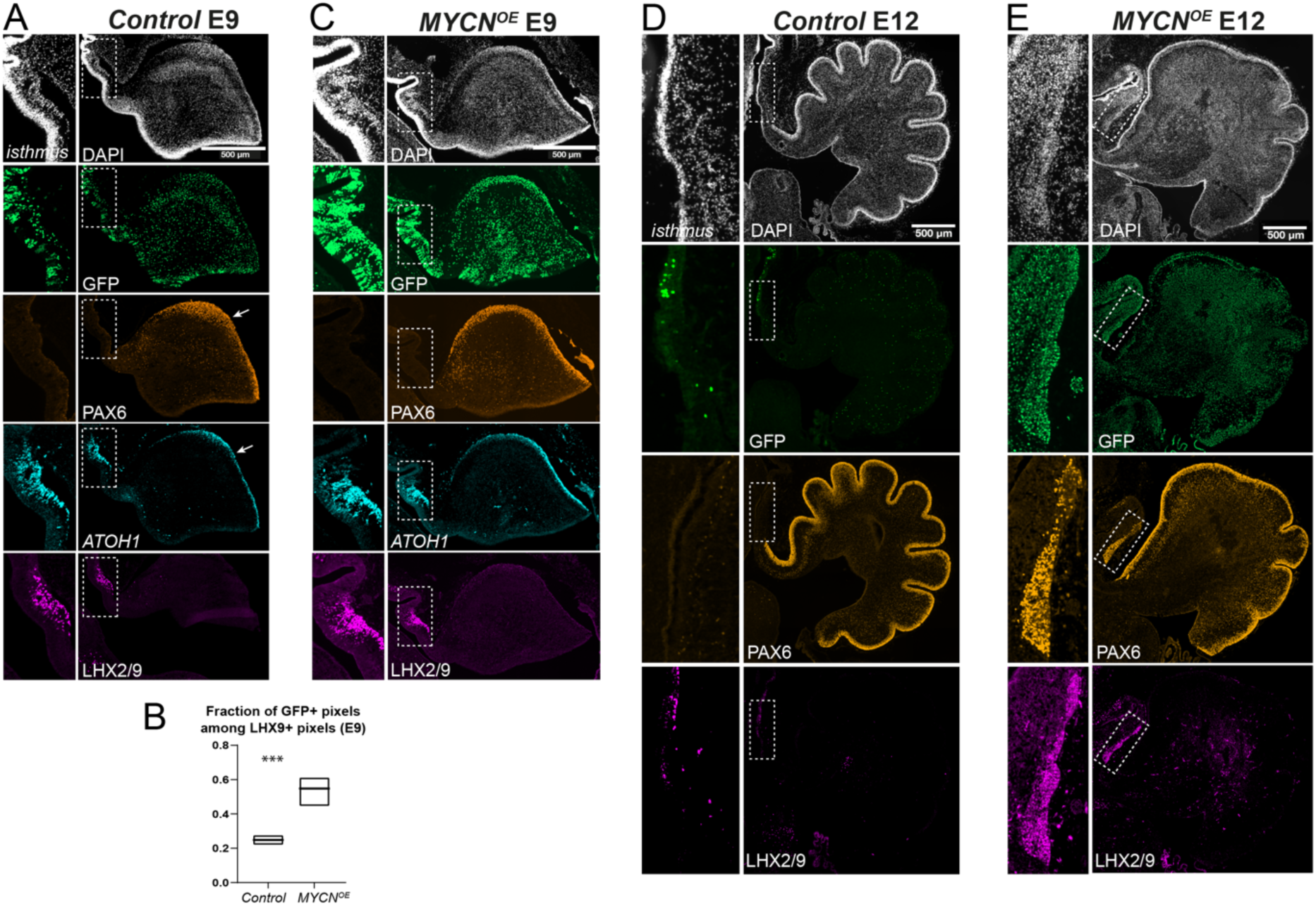
MYCN overexpression expands LHX9^+^ isthmic progenitors and promotes acquisition of a PAX6^+^ GCP-like identity. Representative immunofluorescence images of electroporated embryonic cerebellar/isthmic regions at E9 and E12 (7 and 10 DPE respectively) in control (A and C) and MYCN-overexpressing embryos (B and D). (A) In control embryos at E9, GFP^+^ electroporated cells are detected in the cerebellum and isthmic region (dashed rectangle enlarged on the left side). PAX6 is restricted to the developing EGL (arrow) while Atoh1 is both expressed in the EGL and in the isthmic region. An antibody able to target both LHX9 and LHX2 revealed that LHX2/9 staining is restricted to the isthmus, overlapping with ATOH1. The ATOH1^+^ LHX2/9^+^ region exhibit sparse GFP^+^ cells. (B) Quantitative colocalization analysis between the LHX9 signal and GFP signal in the control and MYCN^OE^ conditions (n=4). (C) Upon MYCN^OE^, PAX6 remained absent from the isthmus at E9. However, large clones of GFP+ cells are detected showing enhanced proliferation of the ATOH1^+^ LHX2/9^+^ progenitor population. (D) At E12 in control embryos, LHX2/9^+^ are rare in the isthmic region, while PAX6 expression remain restricted to the EGL. (E) Upon MYCN^OE^ at E12, a large GFP^+^ LHX2/9^+^ region is observed in the isthmus, showing expansion of the progenitor pool. These cells also acquired robust PAX6 expression, consistent with the reprogramming toward a GCP-like state. Dashed boxes indicate isthmic regions shown at higher magnification.

By E12 (10DPE), LHX9^+^ cells had become sparse in the isthmus of the control condition, suggesting either downregulation of this marker upon differentiation or migration of their progeny (Figure 5D), while PAX6 remained confined to the EGL (Figure 5D). Strikingly, in the *MYCN^OE^*condition, PAX6 was now robustly upregulated in isthmic progenitors that kept expressing LHX9, consistent with our single- cell data (Figure 5E). Thus, *MYCN* overexpression first induces overproliferation and then *de novo* expression of PAX6 in isthmic progenitors.

Together, these results indicate that the extracerebellar tumor does not originate from preexisting PAX6^+^ ATOH1^+^ LHX9^+^ progenitors. Instead, this work identifies ATOH1^+^ LHX9^+^ isthmic progenitors as the likely cell of origin for the extracerebellar SHH medulloblastoma-like tumor. Moreover, they show that PAX6 expression is acquired several days after initiation of MYCN overexpression and the metabolic reprogramming of ATOH1^+^ LHX9^+^ progenitors. This suggests that MYCN progressively redirect the developmental trajectory of these progenitors toward a hybrid isthmic/granule lineage identity.

### PAX6 transcriptional activity sustains expansion of MYCN-reprogrammed isthmic progenitors

To determine whether the derailing of isthmic progenitors toward a GCP-like state is required for tumor formation, we sought to block PAX6 activity during MYCN overexpression. For this purpose, we co-electroporated a dominant-negative PAX6 construct (PAX6-DN), lacking the transactivation domain, for co-expression with MYCN (Figure 6A). As a control, PAX6-DN construct was electroporated alone. This led to reduced cerebellar size at E15 (13DPE) (Figure 6B) consistent with limited maintenance of granule cells, as already observed in mouse PAX6 mutant chimera ^53^.

**Figure 6.**
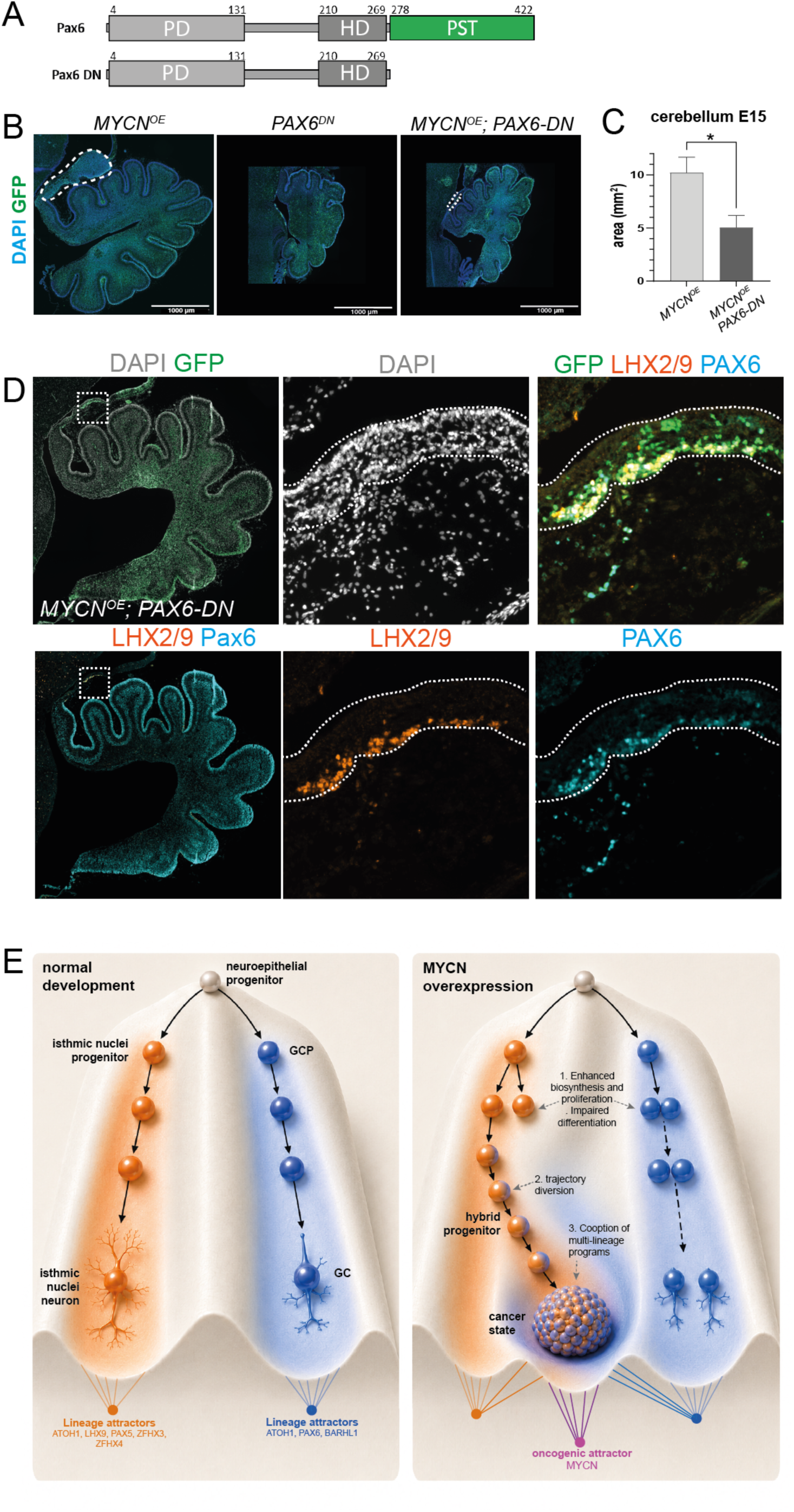
PAX6 transcriptional activity is required to confer tumorigenic potential to the reprogrammed isthmic progenitors. **(A)** Schematic representation of wild-type PAX6 and the dominant-negative PAX6 variant (PAX6DN). PAX6DN retains the paired domain (PD) and homeodomain (HD) DNA-binding domains but lacks the PST transactivation domain. **(B)** Representative immunofluorescence images of sagittal cerebellar sections from control, MYCN-overexpressing (MYCN^OE^), and MYCN^0E^ PAX6DN embryos analyzed at E15, (13 DPE). GFP marks electroporated cells and DAPI labels nuclei. White dashed lines delineate the isthmic region, highlighting the absence of tumor formation upon PAX6DN co-expression. **(C)** Quantification of cerebellar area showing a significant reduction upon combined MYCN PAX6DN overexpression. **(D)** Immunofluorescence analysis of MYCN^OE^; PAX6DN cerebella. Low-magnification views of the cerebellum are shown in the two left panels, with higher-magnification views of the isthmic region shown in the four right panels. Following MYCN + PAX6DN overexpression, the population of LHX9+ and PAX6+ cells remain detectable in the isthmic region but is markedly reduced compared with the MYCN^OE^ condition, indicating that PAX6 transcriptional activity is required to sustain expansion of reprogrammed isthmic progenitors. **(E)** Working model of normal and MYCN-driven cerebellar lineage trajectories represented as a Waddington-like epigenetic landscape. During normal development, neuroepithelial progenitors segregate into an isthmic nuclei lineage and a granule cell lineage, each stabilized by lineage-specific attractors. Upon MYCN overexpression, enhanced anabolic activity and erosion of the landscape leads to increased proliferation and repression of differentiation (1), lineage identity reprogramming and trajectory redirection (2), and cooption of transcriptional programs from other lineages (3). Consequently, isthmic progenitors are diverted toward a hybrid isthmic/granule progenitor state and become trapped in a MYCN-induced stable cancer state, while granule cell differentiation is impaired.

In the MYCN^OE^ PAX6-DN condition, the cerebellum remained smaller (Figure 6B and 6C), but appeared highly electroporated with a high number of GFP^+^ cells throughout. This shows that PAX6-DN does not block the over-proliferative activity of MYCN, at least in lineages in which endogenous PAX6 activity is not required for specification and proliferation. Strikingly, the extracerebellar tumor normally observed following MYCN overexpression was absent in these conditions (N=7), although numerous

GFP^+^ cells remained in the isthmic region. Most of these cells were positive for both LHX9 and endogenous PAX6 (Figure 6D), indicating that the initial reprogramming of LHX9^+^ progenitors toward a PAX6^+^ GCP-like state was still initiated (the antibody used did not detect the truncated form of PAX6). Thus, blocking PAX6 transcriptional activity did not prevent the lineage reprogramming event, but abolished the tumorigenic expansion of reprogrammed isthmic progenitors. This result implies that PAX6-dependent transcription is necessary for expansion of isthmic progenitors.

Thus, our data support a model in which MYCN promotes tumor formation through a three-step temporal sequence: enhancing biosynthesis and proliferation while suppressing differentiation, redirecting LHX9^+^ isthmic progenitors toward a GCP-like fate, and exploiting PAX6-dependent transcriptional programs to sustain a proliferative progenitor state conducive to tumor growth (Figure 6E).

## Discussion

By tracing tumor initiation from its earliest stages in the developing cerebellum and investigating transcriptional perturbations across development, our study reveals that MYCN’s oncogenic properties depend not only on its ability to enhance biosynthesis and impair differentiation, but also on its capacity to progressively redirect lineage trajectories and co-opt regulatory programs from distinct developmental lineages. Together, these findings suggest that derailment of developmental trajectories is a central mechanism of MYCN-driven tumorigenesis, and may represent a broader principle of oncogene-driven transformation. Our findings further suggest that medulloblastoma initiation may involve the acquisition of a new developmental identity, rather than merely the expansion of an early susceptible progenitor population.

### Most cerebellar progenitors are resistant to MYCN-induced oncogenic transformation

Here, we used an *in vivo* mosaic approach combined with single-cell transcriptomics to investigate in unprecedented details how MYCN overexpression affect the lineage trajectory of cerebellar progenitors during the course of development. Our study shows that MYCN overexpression promoted transient overproliferation of progenitor states across cerebellar lineages with delayed or impaired neuronal maturation. This was accompanied with pan-lineage activation of anabolic and translational programs and repression of differentiation programs. However, no tumors were observed within the cerebellum by the end of development, indicating that cerebellar progenitors are largely resistant to the oncogenic effects of MYCN in this embryonic context. This was well illustrated in the granule lineage, where MYCN overexpression impaired terminal GC maturation without blocking the transition from GCPs to immature GCs. These observations challenged our expectation that the GC lineage would be the most susceptible to the oncogenic effects of MYCN. Altogether, these findings are consistent with the established ability of MYC family transcription factors to drive metabolic reprogramming and at least transiently sustain stem cell–like states ^54,55^. Importantly, however, they show that enhancement of anabolic/biosynthetic programs, together with repression of differentiation, is insufficient to initiate tumorigenesis in the context of cerebellar development. Whether this reflects cell-intrinsic resistance to MYCN-driven oncogenic transformation or a non-permissive native microenvironment remains to be investigated.

### MYCN-induced lineage infidelity promotes oncogenesis

Strikingly, MYCN rapidly drove SHH medulloblastoma-like tumors from the isthmus, revealing a restricted progenitor population as particularly susceptible to transformation. We show that these tumors most likely arise from LHX9^+^/ATOH1^+^ progenitors located in the early isthmus, rather than from the rhombic lip or EGL. MYCN initially boosted the proliferation of these progenitors which initially lacked PAX6 expression. Notably, PAX6 became activated over time between E9 and E12, indicating progressive acquisition of granule lineage features. Thus, while it is well-established that medulloblastoma intra-tumoral heterogeneity mirrors normal cerebellar differentiation programs, our finding suggests that this heterogeneity does not necessarily reflects the native lineage of the cell of origin. Instead, oncogenes like MYCN may trigger lineage infidelity, challenging the notion that SHH medulloblastoma necessarily arise from the granule cell lineage.

Importantly, our data indicate that lineage infidelity is not a by-product of transformation but a prerequisite for it, as inhibition of PAX6 transcriptional activity suppressed MYCN-driven overgrowth. These findings therefore support the idea that MYCN oncogenicity in the cerebellum relies not simply on enhanced activation of biosynthetic pathways and repression of differentiation, but also on its ability to erode lineage boundaries in order to redirect lineage identity (Figure 6E). It remains unclear how PAX6, and other lineage-specific features, contribute to trap isthmic progenitors in a cancer state (Figure 6E). It is also unknown why the ability to derail trajectory and cause transformation is only restricted to rare cell states.

Our observation that some LHX9^+^ MYCN-overexpressing progenitors are present in the EGL of anterior lobules at late stages (E19) raises the possibility that they migrate from their primary site to populate the canonical niche of SHH medulloblastoma. Interestingly, this tropism to the EGL is also observed when SHH, but not G3 or G4 medulloblastoma-derived human cell lines are grafted into avian embryos^56^. Although it is unknown whether the ATOH1^+^ LHX9^+^ progenitor pools can be at the origin of medulloblastoma in human, our study raises the possibility that some SHH-medulloblastomas do not originate in the EGL, but colonize the latter during the course of tumorigenesis. Lastly, the long-term reprogramming dynamics of these cells remain unknown. They may progressively lose their initial identity as they grow within the EGL, further obscuring their developmental origin at the time of diagnosis.

While we did not observe granule–isthmic co-expression in the human developmental reference or in chick stages examined (E9–E19), a comprehensive chick single-cell atlas of the isthmus/RL/EGL, or clonal lineage tracing approaches, will be required to exclude rare developmental co-expression events, and formally exclude the possibility that the tumor expands from a rare population of ATOH1^+^ LHX9^+^ PAX6^+^ progenitors.

Beyond medulloblastoma, our work resonates with a recent study showing that MYCN overexpression redirects cone precursor trajectories toward states marked by ectopic expression of multiple retinal lineage markers, some of which are required for transformation into a rare subtype of retinoblastoma^57^. Together, these studies point to lineage infidelity as a potentially recurrent mechanism of MYCN- and more globally oncogene-driven tumorigenesis. By eroding lineage restrictions in some neural progenitors, MYCN may allow cooption of specific gene regulatory networks that act to unleash oncogenic potential in otherwise resistant cells, while masking their true developmental origin at diagnosis.

### MYCN-induced plasticity: beyond granule cells ?

Isthmic tumors were not only composed of GCP- and GC-like cells, but instead displayed marked heterogeneity, including substantial astroglial, oligodendroglial, and immune/microglial compartments. This broad phenotypic diversity raises questions about the origin of these cells. Given our untargeted electroporation approach, it is possible that astroglial cells in the tumor originate from VZ-like progenitors. However, it is also tempting to speculate that MYCN can reprogram cells to a more primitive, multipotent-like state reminiscent of ancestral neuroepithelial cells that possess the potential to give generate both the RL and VZ lineages as shown during early development ^24^, thereby enabling the generation of both neuronal and glial progeny. A last alternative explanation is that GCP-like tumor cells could transdifferentiate into astroglial progenitors. On these lines, tumor-derived astroglial populations have been identified in human SHH medulloblastomas, where they actively support the tumor niche ^58^. More broadly, this model provides a powerful framework to define how MYCN reshapes developmental trajectories, how tumor hierarchies and phenotypic heterogeneity emerge, and how oncogene-induced lineage plasticity contributes to medulloblastoma initiation and evolution.

## Aknowledgements

We are grateful to M-C. Delfini for initial training on the avian embryo during the early stages of the project, L. Telley for discussion and help on human single-cell data, X. Morin for tools and reagents. LB was financed by ANR (ANR-18-CE13-0020), Canceropole PACA (projet Emergence 2020), Ligue contre le cancer (Equipe labelisée EL2019_5 and AAPEAC2020.LCC/MC). PdB was funded by Ligue contre le cancer (AAPEAC2020.LCC/MC). WF was funded by Ligue contre le cancer (Equipe labelisée EL2025_2). GG was funded by Centre South-ROCK (INCa-Cancer_18695). CM was supported by Ligue contre le cancer (Equipe labelisée EL2019_5 and EL2025_2) and by a government grant managed by the Agence Nationale de la Recherche under the France 2030 program, with the reference number ANR-24-EXCI-0001, ANR-24-EXCI-0002, ANR-24-EXCI-0003, ANR-24-EXCI-0004, ANR-24-EXCI-0005.

## Materials and Methods

### Ethical statements

All experiments follow the applicable institutional French (Decree no. 2013-118) and European (Directive 2010/63/EU) guidelines for animal use in research.

### Chicken embryos

Fertilized chicken eggs were obtained from EARL les Bruyères (Dangers, France) and incubated horizontally at 38°C in a humidified incubator. Embryos were staged according to the developmental table of Hamburger and Hamilton (HH) ^59^ or by days of incubation (E).

### Plasmids construction

DNA constructs were designed to constitutively express the gene of interest, *MYCN or PAX6DN*, under the control of a strong ubiquitous CAGGS promoter, with a nuclear or a cytoplasmic GFP downstream respectively, separated by an internal ribosomal entry site (IRES). Constitutive expression was achieved using the Tol2 transposon system, which enables genomic integration of the expression cassette. This was done by co-electroporating the pTol2 construct with a plasmid encoding the Tol2 transposase (pTol2-H2B-GFP and pCAGGS-T2TP; both plasmids were gifts from Dr. Xavier Morin). The coding sequence of *MYCN* and PAX6DN were amplified by PCR from chick cDNA using the following primers:

MYCN-Forward: CTCAAGCTTCGAATTCATGCCGGGAATGATCAGCAAG;

MYCN-Reverse: GTCGACTGCAGAATTCTTAGCAAGTCCGCTTGTACTCTATTTTCT.

PAX6DN-Forward: CTCAAGCTTCGAATTATGCAGAACAGTCACAGCGG

PAX6DN-Reverse: GTCGACTGCAGAATTTTAGATGGGGATGTGGCTGGG

PCR amplifications were performed using PrimeSTAR Max DNA Polymerase (Takara). The PCR product of MYCN was digested with EcoRI and inserted into the MfeI site of the pTol2-IRES-H2B-GFP by ligation. The ClonExpress II One Step Cloning Kit was used to insert PAX6DN PCR product in ptol2-IRES-GFP digested with EcoRI. All the plasmids used for electroporation were purified using a Nucleobond Xtra Midi kit (Macherey-Nagel) and verified by sequencing.

### *In ovo* electroporation

The plasmids used for the gain-of-function experiments consisted of a solution containing pCAGGS-T2TP, pTol2-MYCN-IRES-H2B-GFP and/or pTol2-PAX6DN-IRES-GFP at final concentrations of 0,3 µg/µL and 1,7 µg/µL, respectively. For the control embryos pTol2-IRES-H2B-GFP was used in place of pTol2- MYCN-IRES-H2B-GFP. Fast Green tracer dye was added to the DNA solution for visualization. The eggs were windowed, and the solution was injected into HH8-HH9 chick hindbrains. L-shaped platinum electrodes (BTX-45-0115) were positioned along the neural folds of the midbrain-hindbrain boundary to target the presumptive cerebellum. Electroporation was performed unilaterally using three 50 ms pulses of 18 V at 100 ms intervals, delivered with TSS20 Ovodyne electroporator and EP21 current amplifier (Intra-cel). After electroporation, five drops of PBS solution containing penicillin and streptomycin (Sigma) were administered on top of the yolk before being resealed with transparent tape and returned to the incubator for the desired duration.

### Tissue processing, immunofluorescence, in situ hybridisation and imaging

#### Tissue section

Brains or embryos were fixed in 4% buffered formaldehyde in PBS and then treated with a sucrose gradient (15% and 30% in PBS), embedded in the OCT (optimal cutting temperature) medium, and stored at -80°C. OCT blocks were subsequently sectioned into 12 µm slices using a Leica cryostat and the slides were either stored at -20°C or used directly for FISH and/or immunofluorescence.

#### Immunofluorescence

The slides were rehydrated in PBS and then blocked with 10% goat serum, 3% BSA, 0.4% Triton X-100 in PBS for one hour. The primary antibodies were incubated overnight diluted in the same solution at 4°C. The following primary antibodies were used in this study: chicken anti-GFP 1:1000 (1020 AVES), rabbit anti-SOX2 1:1000 (ab97959 Abcam), rat anti-pH3 1:2000 (S10, 6570 Millipore), mouse anti-NeuN 1:500 (MAB377, Millipore), mouse anti-HuC/HuD 1:500 (A21271, ThermoFisher), mouse anti-LHX2/9 1:20 (1C11, DSHB), mouse anti-PAX6 1:100 (DSHB), rabbit anti- PAX6 1:1000 (AB2237, Millipore). For LHX2/9 staining, an additional antigen retrieval step was performed by incubating the slides in 0.2X SSC buffer at 65 °C. After blocking, sections were incubated with mouse anti-LHX2/9 1:20, followed by an HRP-conjugated anti-mouse secondary antibody (715035151, JIR) and revealed using a TSA-Plus Cyanin-3 kit (Perkin Elmer, Germany).

Secondary antibodies used for the other primary antibodies were Alexa Fluor-conjugated anti-chicken, anti-rabbit, anti-mouse, or anti-rat antibodies (488, 568, or 647; Life Technologies) diluted 1:500 in blocking solution containing Hoechst (1:1000). Incubation was performed for 1h at room temperature.

Slides were washed in PBS, mounted with Thermo Scientific Shandon Immu-Mount, and imaged using a Zeiss Z1 fluorescence microscope.

Fluorescent in situ hybridization on tissue section: The RNA probe used for in situ hybridization targeted Atoh1. The probe template was generated by PCR amplification from chick genomic DNA using the following primers:

Forward: AATCTACATCAGCGCCCTCG;

Reverse: TAATACGACTCACTATAGGGCCACGAGAGAAGAGTTCGCGT.

Notably, 1 M betaine was required for efficient amplification of Atoh1. The reverse primer contained a T7 promoter sequence to allow for in vitro transcription. The PCR product was purified and used as a template for RNA probe synthesis using a T7 RNA polymerase-based transcription reaction with DIG RNA Labeling Mix (Roche), following the manufacturer’s instructions.

Slides were rehydrated in PBS, then treated with 3% H₂O₂ in PBS for 10 minutes, washed in PBS, and incubated in 0.1 M triethanolamine containing 0.25% acetic anhydride for 10 minutes. Pre-hybridization was performed with a hybridization buffer (50% formamide, 5× SSC, 5× Denhardts, 250 µg/mL yeast tRNA and 500 µg/mL herring sperm DNA) for 3h at room temperature and incubated in the same buffer with DIG-labelled RNA probes overnight at 60 °C in a wet chamber. The slides were then washed twice with 0.2× SSC for 30 min at 65 °C. After 5 min in the TNT buffer (100 mM Tris pH 7.5; 150 mM NaCl; 0.1% Tween-20), they were blocked for 1 h in a buffer containing 1× TNT, 1% blocking reagent (Roche) and 10% goat serum, then incubated in the same buffer for 3 h with anti-DIG-POD antibodies (1:500, Roche) and revealed using a TSA-Plus Cyanin-3 kit (Perkin Elmer, Germany). The slides can then proceed to immunofluorescence protocol.

### Histological quantification

Image analysis and signal quantification of cerebellar sections was performed using the open-source software platform QuPath (v0.7.0) ^60^. Cerebellar areas were defined by manual annotation of regions of interest (ROIs) on sagittal sections, and the total area of each ROI was measured using the area measurement tool. To quantify GFP positive cells, nuclei were first detected using the cell detection function applied on the DAPI channel. Detected cells were subsequently classified as GFP-positive based on GFP channel fluorescence intensity thresholding. The proportion of GFP-positive cells among all DAPI-detected nuclei was calculated for each section. At least three embryos per condition and at least three sagittal sections per embryo were analyzed. All plots and statistical analyses were performed in RStudio.

### Mitotic Density Mapping

Mitotic-cell density maps were generated from PH3 mask images and overlaid onto the corresponding DAPI-stained cerebellar section images. PH3-positive mitotic cells were identified from a binary PH3 mask image, in which the minority pixel value represented segmented PH3-positive objects. Connected PH3-positive components were detected and filtered by object size to retain mitotic-cell-sized objects only. Objects with an area between 4 and 100 pixels were included in the analysis (spot_area_median: 10.00 pixels).

For each retained PH3-positive object, the centroid position was calculated and used as a mitotic-cell coordinate. These coordinates were binned onto a regular spatial grid with a 48-pixel grid size and converted into a local mitotic-count map. To visualize regional mitotic density, the count map was smoothed using a Gaussian kernel with a 160-pixel smoothing radius, generating local density domains while preserving broader spatial clustering patterns.

The smoothed density map was normalized to the 99th percentile of positive density values to reduce the influence of extreme local maxima. Density values were then assigned to discrete color classes ranging from very low to highest density. The resulting color-coded density map was overlaid semi-transparently onto the matched DAPI image, used as the anatomical background.

Image processing was performed using a custom Python script. Quality control images showing detected PH3-positive objects over the DAPI background were generated to verify that the density map reflected PH3-positive mitotic-cell distribution rather than background signal.

### Sc-RNA-seq of the chick cerebellum and isthmic tumor

Tissue Dissection and Dissociation for Single-Cell RNA Sequencing: three sample types were collected at embryonic day 15 (E15, 13 days post-electroporation): control posterior lobe (GFP-electroporated), MYCN^OE^ posterior lobe, and extra-cerebellar tumor. For control and MYCN^OE^ posterior lobe samples, the three most posterior cerebellar lobules were microdissected after screening for GFP fluorescence using a LUMAR V12 stereomicroscope (Zeiss).

Tissue dissociation was then performed using 0.125% Trypsin-EDTA in HBSS (without Ca²⁺ and Mg²⁺). Samples were triturated and incubated for 15 min at 37°C with agitation (500 rpm). After incubation, tissue was disrupted using a 21G needle and neutralized with DMEM/F12 + 10% FBS. The dissociated cells were filtered through a 30 µm MACS SmartStrainer (Miltenyi) and resuspended in Neurobasal Medium. The cell suspension was kept on ice and processed for FACS sorting and single-cell RNA sequencing.

FACS sorting and Single cell RNA Sequencing: Chicken cell supension was stained with DAPI (Life Technologies) as a viability marker. Live cells were sorted on BD FACS Aria III SORP (BD Biosciences), centrifuged and resuspended in PBS - 0.04 %(m/w) BSA. For single cell RNA-seq experiment, sorted cells were counted on Attune NxT flow cytometer (Thermo Fisher Scientific). 20,000 cells were encapsulated using chip G on Chromium Controller (10X Genomics) and processed with Chromium Single Cell 3’ Reagent Kit (v3.1) to generate cDNA libraries. Sequencing of libraries was performed on NovaSeq X (Illumina) by Macrogen (Amsterdam, Netherlands).

Custom reference genome preparation and single-cell RNA-seq alignment

The Ensembl *Gallus gallus* reference genome (GRCg7b, release 109) was used as the base for alignment. The original Ensembl annotation for Atoh1 (gene ID: ENSGALG00010008510) contained only the coding sequence (CDS: 36,619,092–36,619,913). Two additional features were manually added to the GTF file: the 5’ UTR region (36,618,981–36,619,091) and the 3’ UTR region (36,619,914–36,620,518). A GFP coding sequence (720 bp) was added as an independent contig (chrGFP) with corresponding annotation. The custom reference genome was indexed using CellRanger mkref, and sequencing libraries were processed with CellRanger count (v8.0.1) using default parameters for alignment, barcode filtering, and gene quantification.

### Cell cycle analysis

The Seurat package provided cell cycle-related genes. The original list contained 43 S-phase markers and 54 G2/M-phase markers which were converted to Gallus gallus orthologs via gProfiler (Ensembl v109) and utilized to determine each cell’s stage by executing the CellCycleScoring function within Seurat.

### PCA, UMAP clustering

CellRanger outputs were processed using Seurat (v4.3.0) in R (v4.1.3). Cells were filtered (1,000–7,000 genes, 1,000–40,000 UMIs, <15% mitochondrial, <20% hemoglobin) based on 38 curated chicken mitochondrial genes (available in the GitHub repository). Data were log-normalized, the 2,000 most variable genes were identified, and cell cycle effects were regressed out during scaling. PCA was performed and 30–40 PCs (dataset-dependent) were used for UMAP and SNN graph-based clustering. Cluster resolution was adjusted per dataset.

### Cell Type Annotation

Cell types were annotated using a label transfer approach with a human reference dataset ^17^ via Seurat’s FindTransferAnchors and TransferData functions. Module scores for reference marker genes were computed to refine cluster identities. Manual annotation was performed based on results obtained after these two different approaches and known markers.

### Differential gene expression, Venn diagrams and gene ontology analysis

Differentially expressed genes (DEGs) were identified using Seurat’s FindAllMarkers and FindMarkers functions (test used “wilcox, “min.pct” set to 0.25 and log fold-change cutoff set to 0.25). To investigate transcriptional similarities and differences among the cell populations identified by scRNA-seq, the 200 most upregulated and the 200 most downregulated DEGs comparing GFP-positive to GFP-negative cells within each cell type were selected to construct Venn diagrams using the R package VennDiagram (v1.7.3). This approach enabled the identification of both shared and cell type-specific regulated genes. Gene ontology (GO) enrichment analysis was subsequently performed with ShinyGO (v0.80; http://bioinformatics.sdstate.edu/go/), using Gallus gallus as the reference species, the GO Biological Process database, and a false discovery rate (FDR) threshold set at 0.05 ^61^.

### Dataset integration and batch correction

To enable comparative analysis across experimental conditions, pre-processed single-cell datasets from control posterior lobe, MYCN^OE^ posterior lobe, and extra-cerebellar tumor were integrated using two complementary approaches.

### Simple merging

For some comparative analyses, preprocessed datasets were merged without integration to preserve dataset-specific variability. The merged object retained original count matrices and cluster annotations from individual analyses, allowing direct comparison of cell type distributions across conditions.

### Seurat rPCA-based integration

For comprehensive multi-sample integration, datasets were integrated using Seurat’s reciprocal PCA (rPCA) workflow ^62^. Integration anchors were identified using the 2,000 most variable features shared across datasets (FindIntegrationAnchors function with reduction = "rpca"). The datasets were then integrated using canonical correlation analysis (CCA)-based integration (IntegrateData function), which creates a batch-corrected "integrated" assay while preserving original count data in the "RNA" assay. The integrated expression matrix was scaled, and PCA was performed on variable features. Based on elbow plot inspection, the first 40 principal components were used for UMAP dimensionality reduction and shared nearest-neighbor (SNN) graph-based clustering with a resolution parameter of 2.5.

Both approaches were used depending on the analytical goal: integration for identifying shared cell populations across conditions, and merging for preserving condition-specific signatures and comparing cluster compositions.

### Integration of extracerebellar tumor-derived pseudo-bulk profiles with pediatric brain tumor reference datasets

Pseudo-bulk generation: Preprocessed extra-cerebellar single-cell RNA-seq data were subset into three major cell lineages: astroglia, granule cells, and oligodendrocytes. Cells from each lineage were randomly partitioned into three balanced groups. Gene expression counts were aggregated per group to generate pseudo-bulk samples, filtered to retain genes with total counts > 5. The resulting matrix was normalized using Variance Stabilizing Transformation (VST) and subjected to Principal Component Analysis (PCA).

Integration with Sturm dataset (GSE73038) ^50^: Microarray data (GSE73038) were retrieved using GEOquery, filtered (counts > 5), and annotated according to tumor subtype. Affymetrix probe identifiers were mapped to human gene symbols using g:Profiler. Pseudo-bulk and Sturm datasets were merged after ensuring gene and sample alignment between matrices.

Batch correction and integrative analysis: Batch effects between datasets were corrected using ComBat (sva). Uniform Manifold Approximation and Projection (UMAP) (n_neighbors = 15, min_dist = 0.1) was performed on corrected data, followed by Louvain clustering (resolution = 1) to identify cell subgroups. Results were visualized with confidence ellipses.

### Trajectory analysis

Spliced and unspliced counts were generated using velocyto (v0.17.17) ^63^. Loom files from the three conditions were merged and analyzed with scVelo (v0.2.4). Data were filtered (min_shared_counts=20, n_top_genes=2000) and moments calculated (npcs=30, n_neighbors=30). RNA velocity was estimated using the stochastic model, refined with dynamical modeling (scv.tl.recover_dynamics()), and latent time was inferred. UMAP coordinates and cluster annotations from Seurat were transferred for visualization consistency. Velocity vectors were displayed as streamlines on UMAP embeddings.

### Statistical analysis and visualization

All statistical analyses were performed in R (v4.1.3). All figures were generated using ggplot2 (v3.4.4), either directly or through Seurat’s plotting functions.

### Single-cell transcriptome analysis of the human developing cerebellum

Human cerebellar single-nucleus RNA-sequencing data were obtained from the SingleCellExperiment object generated by Sepp and colleagues ^17^. Nuclei from 7, 8, 9, 11, 17 and 20 weeks post-conception were retained, and related cell-state annotations were grouped into broader populations. UMI counts were normalized in Seurat using the LogNormalize method with a scale factor of 10,000. For each gene and cell population, mean normalized expression and the proportion of nuclei with non-zero expression were calculated. Expression values were scaled across populations for each gene and limited to the range −2.5 to 2.5. Genes and cell populations were hierarchically clustered using Euclidean distance and the Ward.D2 method.

## Data and code availability

Raw and processed single-cell RNA-seq data generated in this study are publicly available on ArrayExpress under accession number E-MTAB-17146. Analysis scripts (R Markdown) are deposited on GitHub: https://github.com/MaurangeLab/Bouteille-et-al.2026.

## Declaration of generative AI and AI-assisted technologies in the manuscript preparation process

Generative AI-assisted tools were used to support manuscript and figure preparation. ChatGPT was used to assist with editing and polishing the manuscript text for clarity and readability. For the mitotic density map (Figure 1F), ChatGPT assistance was used to develop and run a custom image-analysis workflow in Python. The detected PH3-positive objects and resulting density maps were manually reviewed for consistency with the source images. ChatGPT was also used to assist in the design and preparation of the schematic illustration in Figure 6E. The final scheme was reviewed and edited by the authors to ensure scientific accuracy and consistency with the conclusions of the study. The authors take full responsibility for the accuracy, interpretation, and final presentation of all AI-assisted outputs.

## References

1. Stiles, J., and Jernigan, T.L. (2010). The Basics of Brain Development. Neuropsychol Rev 20, 327–348. 10.1007/s11065-010-9148-4.

2. Taverna, E., Götz, M., and Huttner, W.B. (2014). The Cell Biology of Neurogenesis: Toward an Understanding of the Development and Evolution of the Neocortex. Annual Review of Cell and Developmental Biology 30, 465–502. 10.1146/annurev-cellbio-101011-155801.

3. Wang, L., Wang, C., Moriano, J.A., Chen, S., Zuo, G., Cebrián-Silla, A., Zhang, S., Mukhtar, T., Wang, S., Song, M., et al. (2025). Molecular and cellular dynamics of the developing human neocortex. Nature 647, 169–178. 10.1038/s41586-024-08351-7.

4. Chen, X., Yang, W., Roberts, C.W.M., and Zhang, J. (2024). Developmental origins shape the paediatric cancer genome. Nat Rev Cancer 24, 382–398. 10.1038/s41568-024-00684-9.

5. Kiang, K.M., Wong, Y.K.H., Sengupta, S., Roussel, M.F., and Lu, Q.R. (2025). Developmental Origins and Oncogenesis in Medulloblastoma. Annual Review of Neuroscience 48, 85–102. 10.1146/annurev-neuro-112723-061540.

6. Visvader, J.E. (2011). Cells of origin in cancer. Nature 469, 314–322. 10.1038/nature09781.

7. Gibson, P., Tong, Y.A., Robinson, G., Thompson, M.C., Currle, D.S., Eden, C., Kranenburg, T.A., Hogg, T., Poppleton, H., Martin, J., et al. (2010). Subtypes of medulloblastoma have distinct developmental origins. Nature 468, 1095–1099. 10.1038/nature09587.

8. Tao, R., Han, K., Wu, S.C., Friske, J.D., Roussel, M.F., and Northcott, P.A. (2025). Arrested development: the dysfunctional life history of medulloblastoma. Genes Dev 39, 4–17. 10.1101/gad.351936.124.

9. Cavalli, F.M.G., Remke, M., Rampasek, L., Peacock, J., Shih, D.J.H., Luu, B., Garzia, L., Torchia, J., Nor, C., Morrissy, A.S., et al. (2017). Intertumoral Heterogeneity within Medulloblastoma Subgroups. Cancer Cell 31, 737–754.e6. 10.1016/j.ccell.2017.05.005.

10. Northcott, P.A., Korshunov, A., Witt, H., Hielscher, T., Eberhart, C.G., Mack, S., Bouffet, E., Clifford, S.C., Hawkins, C.E., French, P., et al. (2011). Medulloblastoma comprises four distinct molecular variants. J Clin Oncol 29, 1408–1414. 10.1200/JCO.2009.27.4324.

11. Northcott, P.A., Buchhalter, I., Morrissy, A.S., Hovestadt, V., Weischenfeldt, J., Ehrenberger, T., Grobner, S., Segura-Wang, M., Zichner, T., Rudneva, V.A., et al. (2017). The whole-genome landscape of medulloblastoma subtypes. Nature 547, 311–317. 10.1038/nature22973.

12. Taylor, M.D., Northcott, P.A., Korshunov, A., Remke, M., Cho, Y.-J., Clifford, S.C., Eberhart, C.G., Parsons, D.W., Rutkowski, S., Gajjar, A., et al. (2012). Molecular subgroups of medulloblastoma: the current consensus. Acta Neuropathol 123, 465–472. 10.1007/s00401-011-0922-z.

13. Butts, T., and Wingate, R.J.T. (2026). Chapter One - Simply complex – the structure of the cerebellar neuronal lineage. In Cerebellum Development and Disease Current Topics in Developmental Biology., A. Joyner, R. Sillitoe, and J. Li, eds. (Academic Press), pp. 1–31. 10.1016/bs.ctdb.2026.01.012.

14. Butts, J.C., Wu, S.-R., Durham, M.A., Dhindsa, R.S., Revelli, J.-P., Ljungberg, M.C., Saulnier, O., McLaren, M.E., Taylor, M.D., and Zoghbi, H.Y. (2024). A single-cell transcriptomic map of the developing Atoh1 lineage identifies neural fate decisions and neuronal diversity in the hindbrain. Dev. Cell 59, 2171–2188.e7. 10.1016/j.devcel.2024.07.007.

15. Carter, R.A., Bihannic, L., Rosencrance, C., Hadley, J.L., Tong, Y., Phoenix, T.N., Natarajan, S., Easton, J., Northcott, P.A., and Gawad, C. (2018). A Single-Cell Transcriptional Atlas of the Developing Murine Cerebellum. Curr Biol 28, 2910–2920 e2. 10.1016/j.cub.2018.07.062.

16. Khouri-Farah, N., Guo, Q., Morgan, K., Shin, J., and Li, J.Y.H. (2022). Integrated single-cell transcriptomic and epigenetic study of cell state transition and lineage commitment in embryonic mouse cerebellum. Science Advances 8, eabl9156. 10.1126/sciadv.abl9156.

17. Sepp, M., Leiss, K., Murat, F., Okonechnikov, K., Joshi, P., Leushkin, E., Spänig, L., Mbengue, N., Schneider, C., Schmidt, J., et al. (2024). Cellular development and evolution of the mammalian cerebellum. Nature 625, 788–796. 10.1038/s41586-023-06884-x.

18. Wizeman, J.W., Guo, Q., Wilion, E.M., and Li, J.Y. (2019). Specification of diverse cell types during early neurogenesis of the mouse cerebellum. eLife 8, e42388. 10.7554/eLife.42388.

19. Hoshino, M., Nakamura, S., Mori, K., Kawauchi, T., Terao, M., Nishimura, Y.V., Fukuda, A., Fuse, T., Matsuo, N., Sone, M., et al. (2005). Ptf1a, a bHLH Transcriptional Gene, Defines GABAergic Neuronal Fates in Cerebellum. Neuron 47, 201–213. 10.1016/j.neuron.2005.06.007.

20. Chang, C.-H., Zanini, M., Shirvani, H., Cheng, J.-S., Yu, H., Feng, C.-H., Mercier, A.L., Hung, S.-Y., Forget, A., Wang, C.-H., et al. (2019). Atoh1 Controls Primary Cilia Formation to Allow for SHH-Triggered Granule Neuron Progenitor Proliferation. Developmental Cell 48, 184–199.e5. 10.1016/j.devcel.2018.12.017.

21. Green, M.J., Myat, A.M., Emmenegger, B.A., Wechsler-Reya, R.J., Wilson, L.J., and Wingate, R.J.T. (2014). Independently specified Atoh1 domains define novel developmental compartments in rhombomere 1. Development 141, 389–398. 10.1242/dev.099119.

22. Machold, R., and Fishell, G. (2005). Math1 Is Expressed in Temporally Discrete Pools of Cerebellar Rhombic-Lip Neural Progenitors. Neuron 48, 17–24. 10.1016/j.neuron.2005.08.028.

23. Wang, V.Y., Rose, M.F., and Zoghbi, H.Y. (2005). *Math1* Expression Redefines the Rhombic Lip Derivatives and Reveals Novel Lineages within the Brainstem and Cerebellum. Neuron 48, 31–43. 10.1016/j.neuron.2005.08.024.

24. Zhang, T., Liu, T., Mora, N., Guegan, J., Bertrand, M., Contreras, X., Hansen, A.H., Streicher, C., Anderle, M., Danda, N., et al. (2021). Generation of excitatory and inhibitory neurons from common progenitors via Notch signaling in the cerebellum. Cell Reports 35. 10.1016/j.celrep.2021.109208.

25. Rook, V., Haldipur, P., Millen, K.J., Butts, T., and Wingate, R.J. (2024). BMP signalling facilitates transit amplification in the developing chick and human cerebellum. eLife 12. 10.7554/eLife.92942.2.

26. Hovestadt, V., Smith, K.S., Bihannic, L., Filbin, M.G., Shaw, M.L., Baumgartner, A., DeWitt, J.C., Groves, A., Mayr, L., Weisman, H.R., et al. (2019). Resolving medulloblastoma cellular architecture by single-cell genomics. Nature 572, 74–79. 10.1038/s41586-019-1434-6.

27. Jessa, S., Blanchet-Cohen, A., Krug, B., Vladoiu, M., Coutelier, M., Faury, D., Poreau, B., De Jay, N., Hébert, S., Monlong, J., et al. (2019). Stalled developmental programs at the root of pediatric brain tumors. Nature Genetics 51, 1702–1713. 10.1038/s41588-019-0531-7.

28. Vladoiu, M.C., El-Hamamy, I., Donovan, L.K., Farooq, H., Holgado, B.L., Sundaravadanam, Y., Ramaswamy, V., Hendrikse, L.D., Kumar, S., Mack, S.C., et al. (2019). Childhood cerebellar tumours mirror conserved fetal transcriptional programs. Nature 572, 67–73. 10.1038/s41586-019-1158-7.

29. Okonechnikov, K., Joshi, P., Sepp, M., Leiss, K., Sarropoulos, I., Murat, F., Sill, M., Beck, P., Chan, K.C.-H., Korshunov, A., et al. (2023). Mapping pediatric brain tumors to their origins in the developing cerebellum. Neuro Oncol 25, 1895–1909. 10.1093/neuonc/noad124.

30. Flora, A., Klisch, T.J., Schuster, G., and Zoghbi, H.Y. (2009). Deletion of Atoh1 Disrupts Sonic Hedgehog Signaling in the Developing Cerebellum and Prevents Medulloblastoma. Science 326, 1424–1427. 10.1126/science.1181453.

31. Schüller, U., Heine, V.M., Mao, J., Kho, A.T., Dillon, A.K., Han, Y.-G., Huillard, E., Sun, T., Ligon, A.H., Qian, Y., et al. (2008). Acquisition of Granule Neuron Precursor Identity Is a Critical Determinant of Progenitor Cell Competence to Form Shh-Induced Medulloblastoma. Cancer Cell 14, 123–134. 10.1016/j.ccr.2008.07.005.

32. Dang, C.V. (2012). MYC on the Path to Cancer. Cell 149, 22–35. 10.1016/j.cell.2012.03.003.

33. Hatton, B.A., Knoepfler, P.S., Kenney, A.M., Rowitch, D.H., de Alborán, I.M., Olson, J.M., and Eisenman, R.N. (2006). N-myc Is an Essential Downstream Effector of Shh Signaling during both Normal and Neoplastic Cerebellar Growth. Cancer Res 66, 8655–8661. 10.1158/0008-5472.CAN-06-1621.

34. Swartling, F.J., Grimmer, M.R., Hackett, C.S., Northcott, P.A., Fan, Q.-W., Goldenberg, D.D., Lau, J., Masic, S., Nguyen, K., Yakovenko, S., et al. (2010). Pleiotropic role for MYCN in medulloblastoma. Genes Dev 24, 1059–1072. 10.1101/gad.1907510.

35. Kenney, A.M., Cole, M.D., and Rowitch, D.H. (2003). Nmyc upregulation by sonic hedgehog signaling promotes proliferation in developing cerebellar granule neuron precursors. Development 130, 15–28. 10.1242/dev.00182.

36. Knoepfler, P.S., Cheng, P.F., and Eisenman, R.N. (2002). N-myc is essential during neurogenesis for the rapid expansion of progenitor cell populations and the inhibition of neuronal differentiation. Genes Dev. 16, 2699–2712. 10.1101/gad.1021202.

37. Korshunov, A., Remke, M., Kool, M., Hielscher, T., Northcott, P.A., Williamson, D., Pfaff, E., Witt, H., Jones, D.T.W., Ryzhova, M., et al. (2012). Biological and clinical heterogeneity of MYCN-amplified medulloblastoma. Acta Neuropathol 123, 515–527. 10.1007/s00401-011-0918-8.

38. Pfister, S., Remke, M., Benner, A., Mendrzyk, F., Toedt, G., Felsberg, J., Wittmann, A., Devens, F., Gerber, N.U., Joos, S., et al. (2009). Outcome Prediction in Pediatric Medulloblastoma Based on DNA Copy-Number Aberrations of Chromosomes 6q and 17q and the MYC and MYCN Loci. J Clin Oncol 27, 1627–1636. 10.1200/JCO.2008.17.9432.

39. Roussel, M.F., and Robinson, G.W. (2013). Role of MYC in Medulloblastoma. Cold Spring Harb Perspect Med 3, a014308. 10.1101/cshperspect.a014308.

40. Huang, M., Tailor, J., Zhen, Q., Gillmor, A.H., Miller, M.L., Weishaupt, H., Chen, J., Zheng, T., Nash, E.K., McHenry, L.K., et al. (2019). Engineering Genetic Predisposition in Human Neuroepithelial Stem Cells Recapitulates Medulloblastoma Tumorigenesis. Cell Stem Cell 25, 433–446.e7. 10.1016/j.stem.2019.05.013.

41. Kawauchi, D., Robinson, G., Uziel, T., Gibson, P., Rehg, J., Gao, C., Finkelstein, D., Qu, C., Pounds, S., Ellison, D.W., et al. (2012). A Mouse Model of the Most Aggressive Subgroup of Human Medulloblastoma. Cancer Cell 21, 168–180. 10.1016/j.ccr.2011.12.023.

42. Pöschl, J., Stark, S., Neumann, P., Gröbner, S., Kawauchi, D., Jones, D.T.W., Northcott, P.A., Lichter, P., Pfister, S.M., Kool, M., et al. (2014). Genomic and transcriptomic analyses match medulloblastoma mouse models to their human counterparts. Acta Neuropathol 128, 123–136. 10.1007/s00401-014-1297-8.

43. Swartling, F.J., Savov, V., Persson, A.I., Chen, J., Hackett, C.S., Northcott, P.A., Grimmer, M.R., Lau, J., Chesler, L., Perry, A., et al. (2012). Distinct Neural Stem Cell Populations Give Rise to Disparate Brain Tumors in Response to N-MYC. Cancer Cell 21, 601–613. 10.1016/j.ccr.2012.04.012.

44. Zindy, F., Uziel, T., Ayrault, O., Calabrese, C., Valentine, M., Rehg, J.E., Gilbertson, R.J., Sherr, C.J., and Roussel, M.F. (2007). Genetic Alterations in Mouse Medulloblastomas and Generation of Tumors De novo from Primary Cerebellar Granule Neuron Precursors. Cancer Res 67, 2676–2684. 10.1158/0008-5472.CAN-06-3418.

45. Hill, R.M., Kuijper, S., Lindsey, J.C., Petrie, K., Schwalbe, E.C., Barker, K., Boult, J.K.R., Williamson, D., Ahmad, Z., Hallsworth, A., et al. (2015). Combined MYC and P53 Defects Emerge at Medulloblastoma Relapse and Define Rapidly Progressive, Therapeutically Targetable Disease. Cancer Cell 27, 72–84. 10.1016/j.ccell.2014.11.002.

46. Haldipur, P., Aldinger, K.A., Bernardo, S., Deng, M., Timms, A.E., Overman, L.M., Winter, C., Lisgo, S.N., Razavi, Silvestri, E., et al. (2019). Spatiotemporal expansion of primary progenitor zones in the developing human cerebellum. Science 366, 454–460. 10.1126/science.aax7526.

47. Englund, C., Kowalczyk, T., Daza, R.A.M., Dagan, A., Lau, C., Rose, M.F., and Hevner, R.F. (2006). Unipolar Brush Cells of the Cerebellum Are Produced in the Rhombic Lip and Migrate through Developing White Matter. J. Neurosci. 26, 9184–9195. 10.1523/JNEUROSCI.1610-06.2006.

48. Seto, Y., Nakatani, T., Masuyama, N., Taya, S., Kumai, M., Minaki, Y., Hamaguchi, A., Inoue, Y.U., Inoue, T., Miyashita, S., et al. (2014). Temporal identity transition from Purkinje cell progenitors to GABAergic interneuron progenitors in the cerebellum. Nat Commun 5, 3337. 10.1038/ncomms4337.

49. Kebschull, J.M., Casoni, F., Consalez, G.G., Goldowitz, D., Hawkes, R., Ruigrok, T.J.H., Schilling, K., Wingate, R., Wu, J., Yeung, J., et al. (2024). Cerebellum Lecture: the Cerebellar Nuclei—Core of the Cerebellum. Cerebellum 23, 620–677. 10.1007/s12311-022-01506-0.

50. Sturm, D., Orr, B.A., Toprak, U.H., Hovestadt, V., Jones, D.T.W., Capper, D., Sill, M., Buchhalter, I., Northcott, P.A., Leis, I., et al. (2016). New Brain Tumor Entities Emerge from Molecular Classification of CNS-PNETs. Cell 164, 1060–1072. 10.1016/j.cell.2016.01.015.

51. Funahashi, J., Okafuji, T., Ohuchi, H., Noji, S., Tanaka, H., and Nakamura, H. (1999). Role of Pax-5 in the regulation of a mid-hindbrain organizer’s activity. Development, Growth & Differentiation 41, 59–72. 10.1046/j.1440-169x.1999.00401.x.

52. Butts, T., Green, M.J., and Wingate, R.J.T. (2014). Development of the cerebellum: simple steps to make a ‘little brain.’ Development 141, 4031–4041. 10.1242/dev.106559.

53. Swanson, D.J., and Goldowitz, D. (2011). Experimental Sey mouse chimeras reveal the developmental deficiencies of Pax6-null granule cells in the postnatal cerebellum. Developmental Biology 351, 1–12. 10.1016/j.ydbio.2010.11.018.

54. Poli, V., Fagnocchi, L., Fasciani, A., Cherubini, A., Mazzoleni, S., Ferrillo, S., Miluzio, A., Gaudioso, G., Vaira, V., Turdo, A., et al. (2018). MYC-driven epigenetic reprogramming favors the onset of tumorigenesis by inducing a stem cell-like state. Nat Commun 9, 1024. 10.1038/s41467-018-03264-2.

55. Tjaden, B., Baum, K., Marquardt, V., Simon, M., Trajkovic-Arsic, M., Kouril, T., Siebers, B., Lisec, J., Siveke, J.T., Schulte, J.H., et al. (2020). N-Myc-induced metabolic rewiring creates novel therapeutic vulnerabilities in neuroblastoma. Sci Rep 10, 7157. 10.1038/s41598-020-64040-1.

56. Mallet, M., Martin, F., Thoinet, K., Imbert, C., Sarhadi, M., Ganofsky, J., Diaz, J.T., Fenouil, T., Delloye-Bourgeois, C., Falk, J., et al. (2025). Lineage of origin-specific developmental programs drive the behaviors of malignant cells in an avian embryo model of human Medulloblastoma subgroups. Preprint at bioRxiv, 10.1101/2025.10.09.681440 10.1101/2025.10.09.681440.

57. Singh, H.P., Shayler, D.W.H., Fernandez, G.E., Thornton, M.E., Craft, C.M., Grubbs, B.H., and Cobrinik, D. (2022). An immature, dedifferentiated, and lineage-deconstrained cone precursor origin of N-Myc–initiated retinoblastoma. Proceedings of the National Academy of Sciences 119, e2200721119. 10.1073/pnas.2200721119.

58. Yao, M., Ventura, P.B., Jiang, Y., Rodriguez, F.J., Wang, L., Perry, J.S.A., Yang, Y., Wahl, K., Crittenden, R.B., Bennett, M.L., et al. (2020). Astrocytic trans-Differentiation Completes a Multicellular Paracrine Feedback Loop Required for Medulloblastoma Tumor Growth. Cell 180, 502–520.e19. 10.1016/j.cell.2019.12.024.

59. Hamburger, V., and Hamilton, H.L. (1992). A series of normal stages in the development of the chick embryo. Developmental Dynamics 195, 231–272. 10.1002/aja.1001950404.

60. Bankhead, P., Loughrey, M.B., Fernández, J.A., Dombrowski, Y., McArt, D.G., Dunne, P.D., McQuaid, S., Gray, R.T., Murray, L.J., Coleman, H.G., et al. (2017). QuPath: Open source software for digital pathology image analysis. Sci Rep 7, 16878. 10.1038/s41598-017-17204-5.

61. Ge, S.X., Jung, D., and Yao, R. (2020). ShinyGO: a graphical gene-set enrichment tool for animals and plants. Bioinformatics 36, 2628–2629. 10.1093/bioinformatics/btz931.

62. Hao, Y., Stuart, T., Kowalski, M.H., Choudhary, S., Hoffman, P., Hartman, A., Srivastava, A., Molla, G., Madad, S., Fernandez-Granda, C., et al. (2024). Dictionary learning for integrative, multimodal and scalable single-cell analysis. Nat Biotechnol 42, 293–304. 10.1038/s41587-023-01767-y.

63. La Manno, G., Soldatov, R., Zeisel, A., Braun, E., Hochgerner, H., Petukhov, V., Lidschreiber, K., Kastriti, M.E., Lönnerberg, P., Furlan, A., et al. (2018). RNA velocity of single cells. Nature 560, 494–498. 10.1038/s41586-018-0414-6.

