## Supplementary Figures 1-5 for "MYCN-induced lineage infidelity initiates SHH medulloblastoma-like tumors outside the canonical lineage of origin"

### Supplementary Material

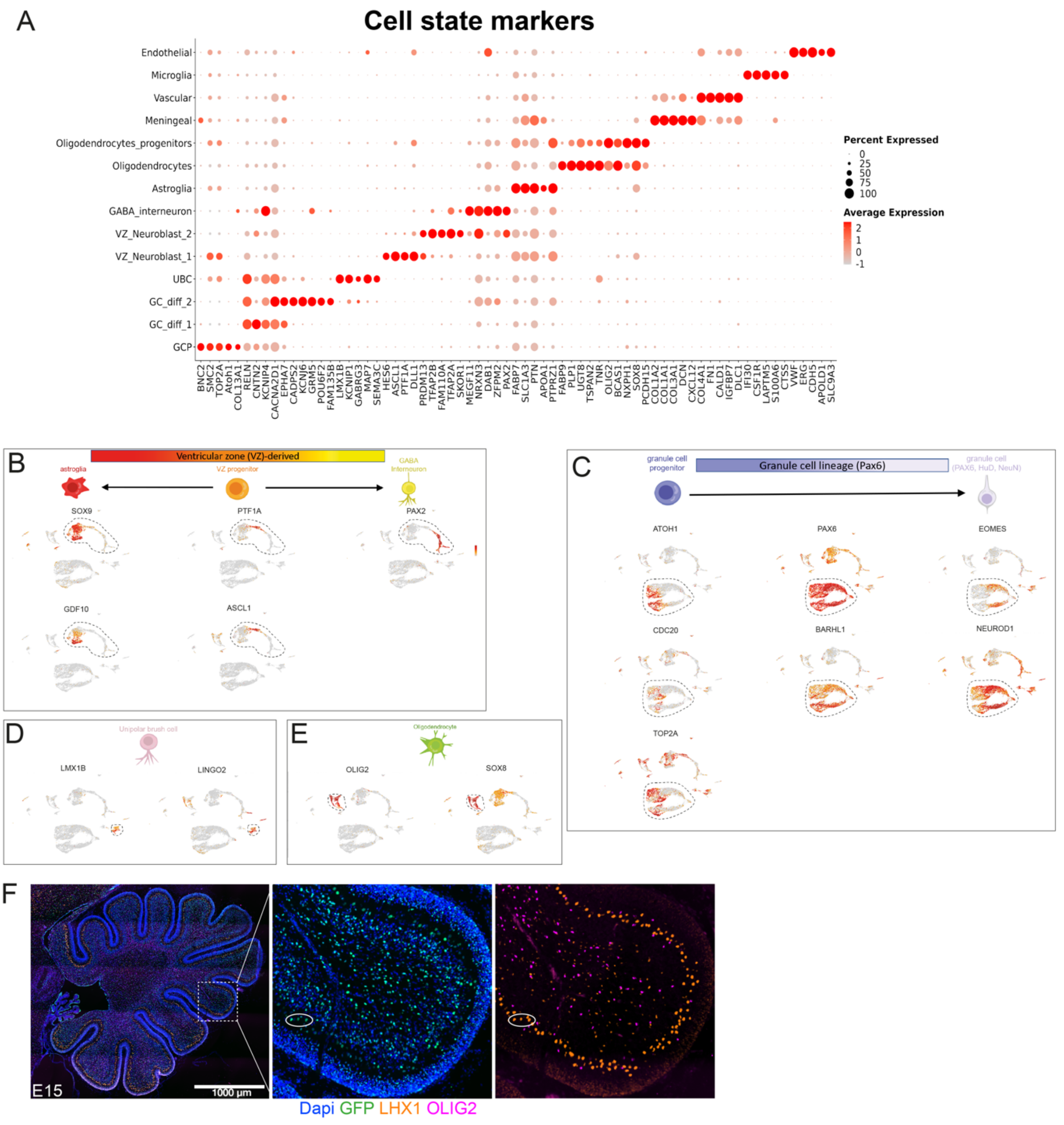


**Figure S1: Chick cerebellar cell types identified by single-cell RNA-seq**

(A) Dot plot showing top differentially expressed genes for annotated cerebellar cell populations. Dot size represents the percentage of cells expressing each gene and color intensity indicates average expression levels. The list of Top differentially-expressed genes are provided in Table S1.
(B) Feature plots depicting expression of various markers for VZ-derived cell states or cell types.
(C) Feature plots depicting expression of various markers for the GC lineage.
(D) Feature plots depicting expression of various markers for presumptive UBCs.
(E) Feature plots depicting expression of various oligodendrocyte markers.
(F) Immunofluorescence images of a sagittal cerebellar section at E15 stained with antibodies labeling the nucleus of Purkinje cells (LHX1) and oligodendrocytes (Olig2). Note that Purkinje cells exhibit a larger nucleus than other cell types and can be efficiently electroporated as denoted by GFP+ nuclei in the white ellipse.

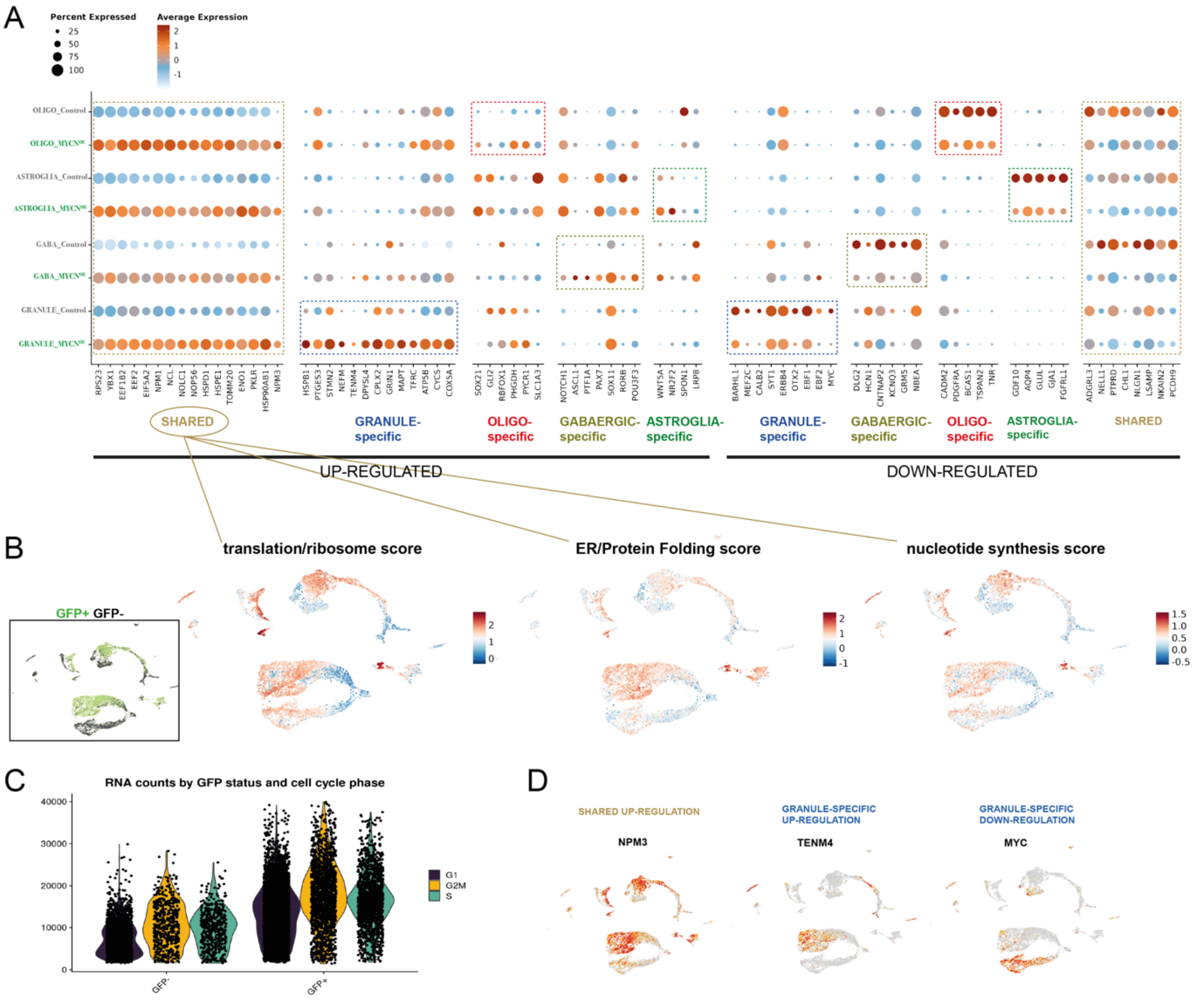


**Figure S2: MYCN-induced transcriptional effects in cerebellar lineages**

(A) Dotplot depicting examples of genes that are up-regulated or down-regulated across cerebellar lineages in a shared or lineage-specific manner.

(B) Feature plots depicting $GFP^+$ and $GFP^-$ cells and module scores for biological processes enriched in the shared MYCN-induced program.

(C) $GFP^+$ cells exhibit a higher content of mRNAs than $GFP^-$ cells independently of the cell cycle phase.

(D) Feature plots exhibiting pan-lineage up-regulation of NPM3 in electroporated cells, GCP-specific up-regulation of TENM4, and granule lineage specific down-regulation of MYC in electroporated cells.

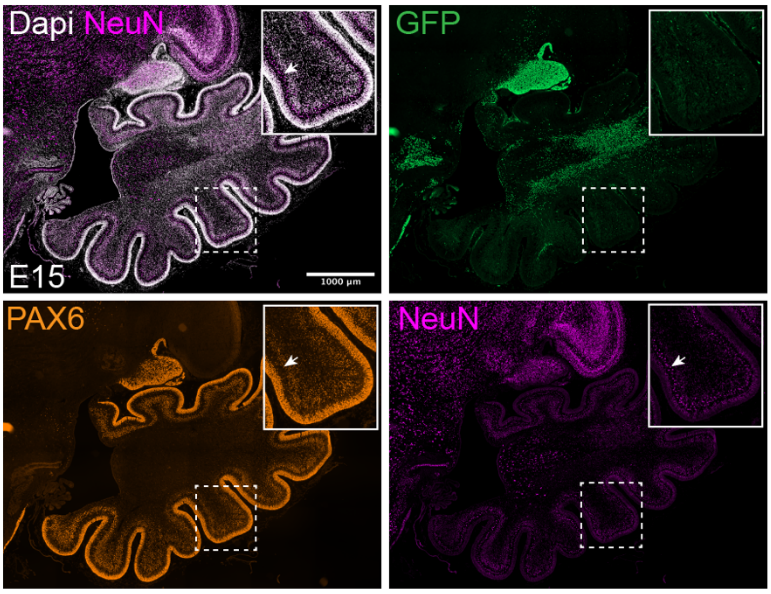


**Figure S3: GCs do not express the differentiation marker NeuN at E15**
PAX6+ GCs do not express NeuN at 15. In contrast, arrow indicates large nuclei of differentiated Purkinje cells that express NeuN but not PAX6.

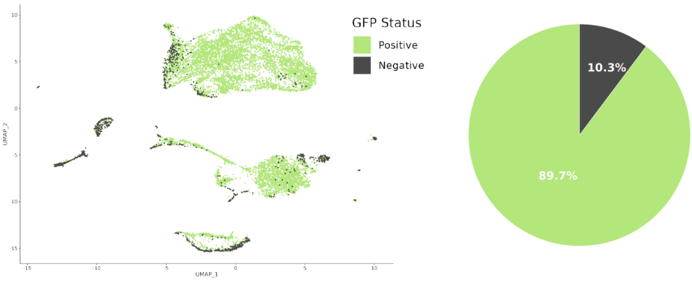


**Figure S4: Analysis of the single-cell data from the extra-cerebellar tumor**
GFP+ and GFP- fractions are depicted in the different clusters. GFP+ cells represent the vast majority in the extracerebellar tumor.

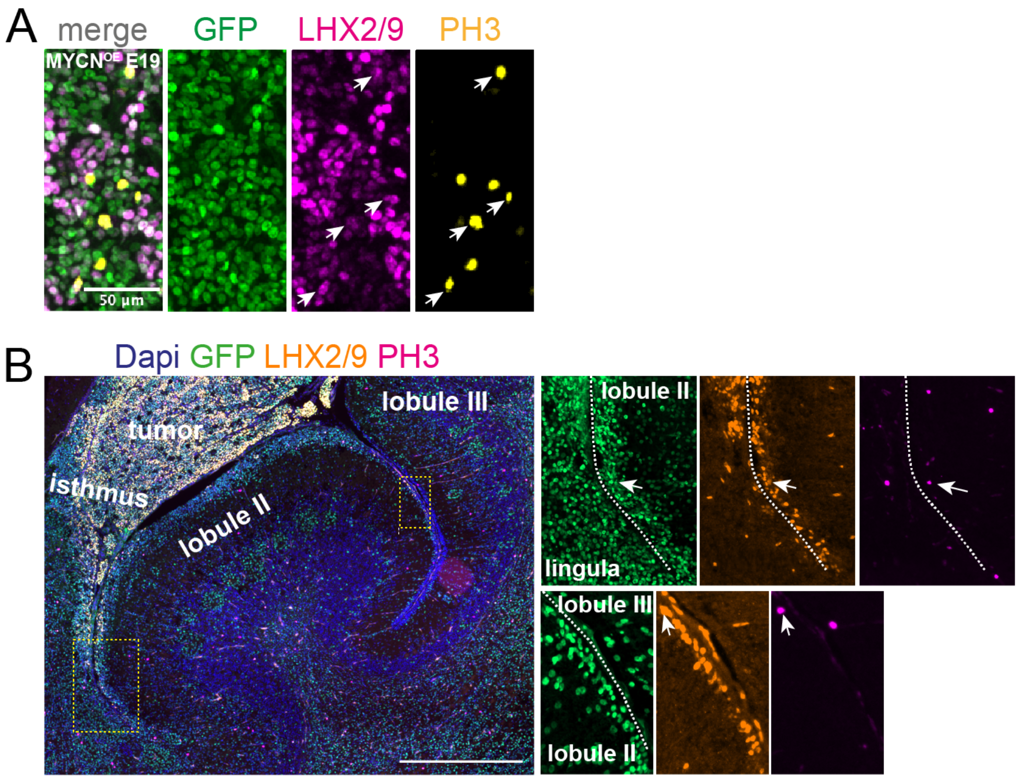


**Figure S5: LHX9+ tumor cells are proliferative and possibly exhibit invasive properties**
(A) High-magnification images of the tumor at E19 showing GFP, LHX2/9, and phospho-histone H3 (PH3) immunostaining. Arrows indicate proliferative LHX2/9$^{+}$ tumor cells. (B) Sagittal section showing GFP, LHX2/9, and PH3 staining in an MYCN-electroporated cerebellum. GFP+ PAX6+ LHX2/9$^{+}$ tumor cells are detected within the EGL of anterior cerebellar lobules, indicating invasion of tumor progenitors into the endogenous GCP niche. Insets show higher magnification of boxed regions.

**Table S1:** Differentially expressed genes across clusters in posterior cerebellar lobules co-electroporated with GFP and MYCN

**Table S2:** Differentially expressed genes between GFP+ and GFP- cells for each lineage

**Table S3:** TOP200 up- and down-regulated genes between GFP+ and GFP- cells for each lineage and Venn intersections

**Table S4:** Top 100 differentially expressed genes between GFP$^{+}$ tumorigenic GCP-like progenitors and GFP$^{+}$ GCPs of the EGL.
